# Segmental duplications and their variation in a complete human genome

**DOI:** 10.1101/2021.05.26.445678

**Authors:** Mitchell R. Vollger, Xavi Guitart, Philip C. Dishuck, Ludovica Mercuri, William T. Harvey, Ariel Gershman, Mark Diekhans, Arvis Sulovari, Katherine M. Munson, Alexandra M. Lewis, Kendra Hoekzema, David Porubsky, Ruiyang Li, Sergey Nurk, Sergey Koren, Karen H. Miga, Adam M. Phillippy, Winston Timp, Mario Ventura, Evan E. Eichler

## Abstract

Despite their importance in disease and evolution, highly identical segmental duplications (SDs) have been among the last regions of the human reference genome (GRCh38) to be finished. Based on a complete telomere-to-telomere human genome (T2T-CHM13), we present the first comprehensive view of human SD organization. SDs account for nearly one-third of the additional sequence increasing the genome-wide estimate from 5.4% to 7.0% (218 Mbp). An analysis of 266 human genomes shows that 91% of the new T2T-CHM13 SD sequence (68.3 Mbp) better represents human copy number. We find that SDs show increased single-nucleotide variation diversity when compared to unique regions; we characterize methylation signatures that correlate with duplicate gene transcription and predict 182 novel protein-coding gene candidates. We find that 63% (35.11/55.7 Mbp) of acrocentric chromosomes consist of SDs distinct from rDNA and satellite sequences. Acrocentric SDs are 1.75-fold longer (p=0.00034) than other SDs, are frequently shared with autosomal pericentromeric regions, and are heteromorphic among human chromosomes. Comparing long-read assemblies from other human (n=12) and nonhuman primate (n=5) genomes, we use the T2T-CHM13 genome to systematically reconstruct the evolution and structural haplotype diversity of biomedically relevant (*LPA, SMN*) and duplicated genes (*TBC1D3, SRGAP2C, ARHGAP11B*) important in the expansion of the human frontal cortex. The analysis reveals unprecedented patterns of structural heterozygosity and massive evolutionary differences in SD organization between humans and their closest living relatives.

## INTRODUCTION

Genomic duplications have long been recognized as important sources of structural change and gene innovation (*1, 2*). In humans, for example, the most recent and highly identical sequences (>90%), referred to as segmental duplications (SDs) (*3*), promote meiotic unequal crossover events contributing to recurrent rearrangements associated with ∼5% of developmental delay and autism (*4*). These same SDs are reservoirs for human-specific genes important in increasing synaptic density and the expansion of the frontal cortex since humans diverged from other ape lineages (*5*–*8*). SDs are also ∼10-fold enriched for normal copy number variation although most of this genetic diversity has yet to be fully characterized or associated with human phenotypes (*9, 10*). Their length (frequently >100 kbp), sequence identity, and extensive structural diversity among human haplotypes have hampered our ability to characterize these regions at a genomic level because sequence reads have been insufficiently long and human haplotypes too structurally diverse to resolve duplicate copies or distinguish allelic variants. One of the first human whole-genome sequence (WGS) assembly drafts based on Sanger sequence reads was almost completely devoid of SDs and their underlying genes (*11, 12*). Similarly, BAC-based approaches to assemble the human genome from different haplotypes led to many misjoins creating *de facto* gaps that took years to resolve (*13*). While combining WGS- and BAC-based data from the first human genomes provided a road map of the SD landscape (*14*), more than 50% of the gaps within the human reference genome have corresponded to regions of complex SDs. The development of genomic resources (*15*–*17*), including BAC libraries and long-read sequence data from complete hydatidiform moles (which represent a single human haplotype), was motivated in large part by efforts to resolve the organization of these regions and concomitantly complete the human reference genome. Using these resources combined with advances in long-read technologies, a gapless human genome assembly (T2T-CHM13) is now complete (*18*). Here, we use this genome assembly to present the most complete view of SDs in a human genome and highlight their importance in advancing our understanding of human genetic diversity, evolution, and disease.

## RESULTS

### SD content and organization

We characterized the SD content of the T2T-CHM13 v1.0 assembly based on sequence read-depth and pairwise sequence alignments (>90% and >1 kbp) (*19*). Our analysis of the assembly identifies 218 Mbp of nonredundant segmentally duplicated sequence within chromosome-level scaffolds, compared to just 167 Mbp in the current reference (GRCh38) (Table 1, Fig. 1). This raises the percent estimate of the human genome that is segmentally duplicated from 5.4% to 7.0%. Five gaps remained in the initial T2T-CHM13 assembly. Each corresponded to a cluster of tandemly repeated rDNA genes on each acrocentric chromosome where we confirm long-read sequence pileups representing the last unresolved SDs of the human genome. To estimate the amount of missing duplicated rDNA sequence, we applied digital droplet PCR (*20*) and a whole-genome Illumina coverage analysis (*18*). Assuming a canonical repeat length of 45 kbp for the rDNA molecule (*21, 22*), we approximate that there are ∼10 Mbp and ∼200 copies of unresolved rDNA sequence (*18*). Including this, the overall SD content of the human genome is 7.0% (6.7% not including rDNA; see Table 1 for statistics breakdown by SD type). These findings are consistent with the subsequent specialized assembly of the rDNA released as part of the T2T-CHM13 v1.1 assembly (Table 1).

**Table 1.** Summary statistics of segmental duplications in T2T CHM13 and GRCh38.

| *Assembly* | *Gbp* | *%*<br>*SD* | *SD*<br>*(Mbp)* | *# SDs* | *inter*<br>*(Mbp)* | *#*<br>*inter* | *intra*<br>*(Mbp)* | *#*<br>*intra* | *acro*<br>*(Mbp)* | *#*<br>*acro* | *peri*<br>*(Mbp)* | *# peri* | *telo*<br>*(Mbp)* | *#*<br>*telo* |
| --- | --- | --- | --- | --- | --- | --- | --- | --- | --- | --- | --- | --- | --- | --- |
| T2T CHM13 | 3.11 | 6.66 | 207.55 | 41285 | 121.10 | 30480 | 142.96 | 10805 | 35.11 | 13264 | 88.61 | 24984 | 10.98 | 4998 |
| GRCh38 | 3.09 | 5.42 | 167.30 | 24280 | 83.56 | 16348 | 120.71 | 7932 | 6.62 | 1407 | 53.94 | 10606 | 8.93 | 1529 |
| Difference | 0.03 | 1.25 | 40.26 | 17005 | 37.54 | 14132 | 22.25 | 2873 | 28.48 | 11857 | 34.66 | 14378 | 2.05 | 3469 |
| New or structurally variable | 0.24 | 33.88 | 81.34 | 25161 | 61.87 | 20579 | 54.93 | 4582 | 35.04 | 13258 | 54.04 | 19607 | 5.62 | 4005 |
| T2T CHM13 v1.1 | 3.11 | 6.99 | 217.54 | 41711 | 131.06 | 30903 | 151.95 | 10808 | 45.07 | 13687 | 95.24 | 25237 | 11.06 | 4997 |

Mbp: the number of non-redundant Mbp of SD; peri: within 5Mbp of the heterochromatin surrounding the centromere; telo: within 500 kbp of the telomere; acro: within the short arms of the acrocentric chromosomes.
Difference: SD content difference between T2T-CHM13 v1.0 and GRCh38.
New or structurally variable: Sequence in T2T-CHM13 that does not have 1 Mbp of synteny with GRCh38 (methods).
GRCh38 contains 149,690,719 of gap sequence included in the reported # of Gbp.
Supplemental note 1.

**Fig. 1.**
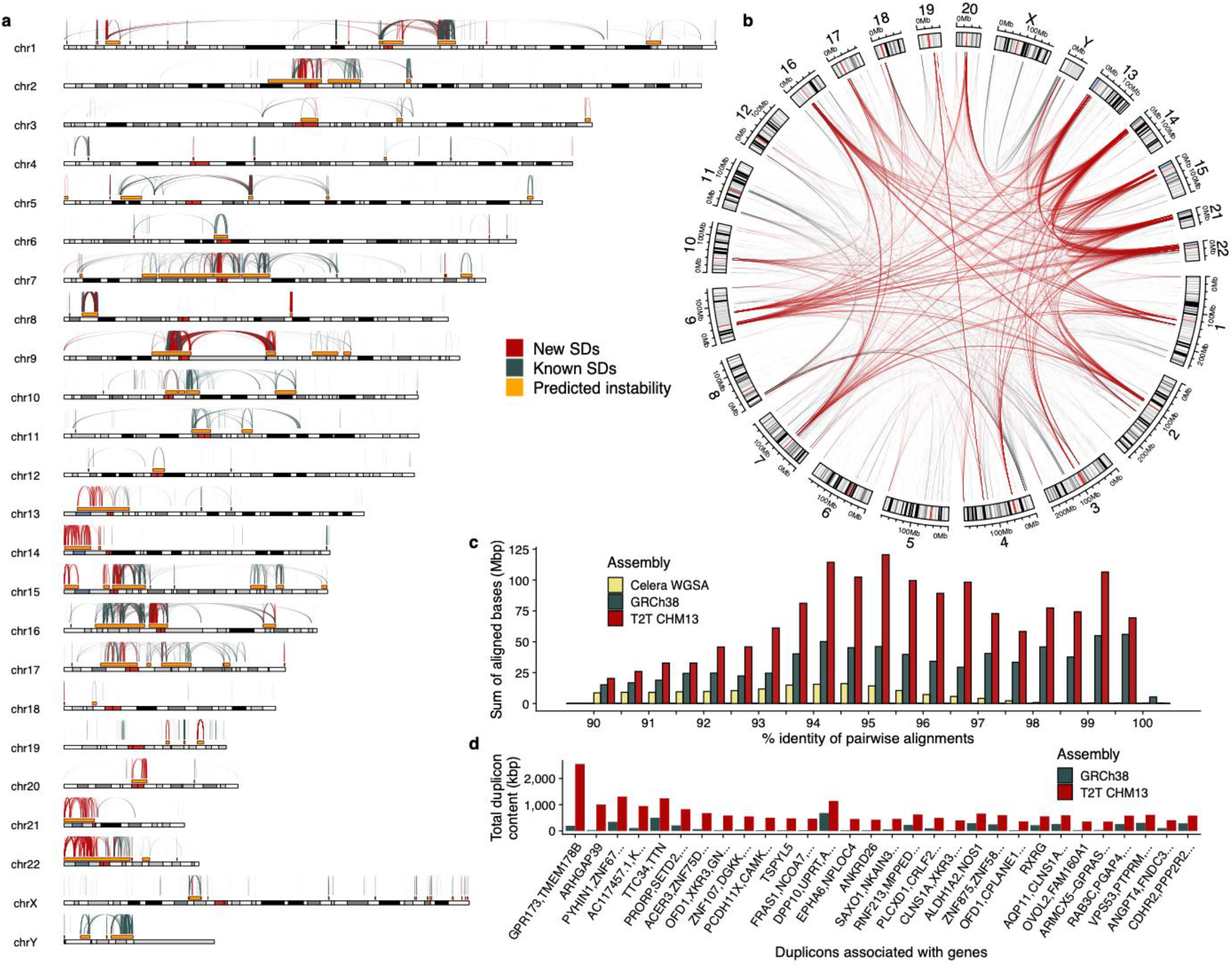
Segmental duplication (SD) content of the T2T-CHM13 genome. **a)** The pattern of novel or structurally variant intrachromosomal duplications in T2T-CHM13 (red) compared to known duplications in GRCh38 (blue-gray). These predict hotspots of genomic instability (gold) flanked by large (>10 kbp), high-identity (>95%) interspersed (>50 kbp) SDs. **b)** Circos plot highlighting novel interchromosomal SDs (red) shows the preponderance of new SDs mapping to pericentromeric and acrocentric regions. **c)** A histogram comparing SD content in different human reference genomes. The sum of bases in pairwise SD alignments stratified by their percent identity for the Celera (yellow, Sanger-based), GRCh38 (blue-gray, BAC-based), and T2T-CHM13 (red, long-read) assemblies. **d)** The 30 genic duplicons (ancestral repeat units) with the greatest copy number difference between GRCh38 and T2T-CHM13 as determined by DupMasker. All of the 30 largest differences are present in T2T-CHM13.

One-third (81.3 Mbp, Supplemental note 1) of SDs are new or differ structurally when comparing the T2T-CHM13 assembly to GRCh38. Most of these involve large, high-identity SDs. For example, there is a 70% increase (41,285/24,280) in the number of SD pairs and a doubling of the number of bases in pairwise alignments with greater than 95% identity (Fig. 1c). Among these new or variable SDs, 13,258 (35.0 Mbp) map to the acrocentric short arms of chromosomes 13, 14, 15, 21, and 22 (Fig. 1b, Table 1), which are assembled for the first time. These SDs do not correspond to rDNA duplications but represent other segments predominantly shared among acrocentric (n=5,332 alignments) and non-acrocentric chromosomes (n= 5,500 alignments, Table S1). In particular, the pericentromeric regions of chromosomes 1, 3, 4, 7, 9, 16 and 20 show the most extensive SD homology with acrocentric DNA (Fig. 1b). Non-rDNA acrocentric SDs are 1.75-fold longer than all other SDs (N50: 74,704 vs. 42,842)--significantly longer than any other defined SD category in the human genome [intrachromosomal, interchromosomal, pericentromeric, and telomeric (Fig. S1)].

We annotated all T2T-CHM13 SDs using DupMasker (*23*), which defines ancestral evolutionary units of duplication based on mammalian outgroups and a repeat graph (*24*). Focusing on duplicons that carry genes or duplicated portions of genes, we identify 30 duplicons that show the greatest copy number change between T2T-CHM13 and GRCh38. These 30 genic SDs represent regions where gene annotation is most likely to change; all predicted differences favor an increase in copy number for the T2T-CHM13 assembly (Fig. 1d, Table S2). We also compared the number of SDs more directly by defining syntenic regions (5 Mbp) between GRCh38 and T2T-CHM13 (Methods). Of the 15 windows with the largest increase, nine mapped to the acrocentric short arms while six were in pericentromeric regions (Fig. S1, Table S3). In particular, the intervals between the centromeric satellite and secondary constrictions (qh regions) on chromosomes 1, 9, and 16 show a 4.6-fold increase in the number of SDs (5,254/1,141) and show the most dramatic differences in organization when compared to GRCh38. SDs in these regions are almost exclusively interchromosomal and depleted for intrachromosomal duplications (Fig. S2-S3).

### Validation and heteromorphic variation

Because the acrocentric short arms as well as the qh regions on chromosomes 1, 9, and 16 were either newly assembled or showed the most significant differences in terms of SD content, we focused first on validating their organization. We mapped available end-sequence data from a human fosmid genome library (*25*) to the T2T-CHM13 assembly and selected nine distinct clones as probes (Fig. 2a) to confirm the patterns of high-identity (>95%) SDs. All 30 of the distinct duplication predictions based on T2T-CHM13 SDs were corroborated by FISH against chromosomal metaphases of the CHM13 cell line (Fig. 2b, Table S4). Interestingly, FISH also revealed nine additional signals not originally predicted by our SD analysis (Fig. S4). However, we were able to identify lower identity duplications confirming seven of these sites leading to an overall concordance of 95% (37/39) between FISH and the T2T-CHM13 SD assembly content. We extended this analysis to five additional human cell lines of diploid origin because both pericentromeric and acrocentric portions of chromosomes have been shown to be cytogenetically heteromorphic (*26*–*28*). In total, we identified 61 distinct cytogenetic locations of which 28 (46%) were fixed while 33 (54%) were variable in their presence or absence on specific homologues (both acrocentric and pericentromeric regions of the human genome) (Fig. S4). Of the 61 FISH signals, all but six were observed in more than one of the six human cell lines indicating that such heteromorphic variation is common and prevalent.

**Fig. 2.**
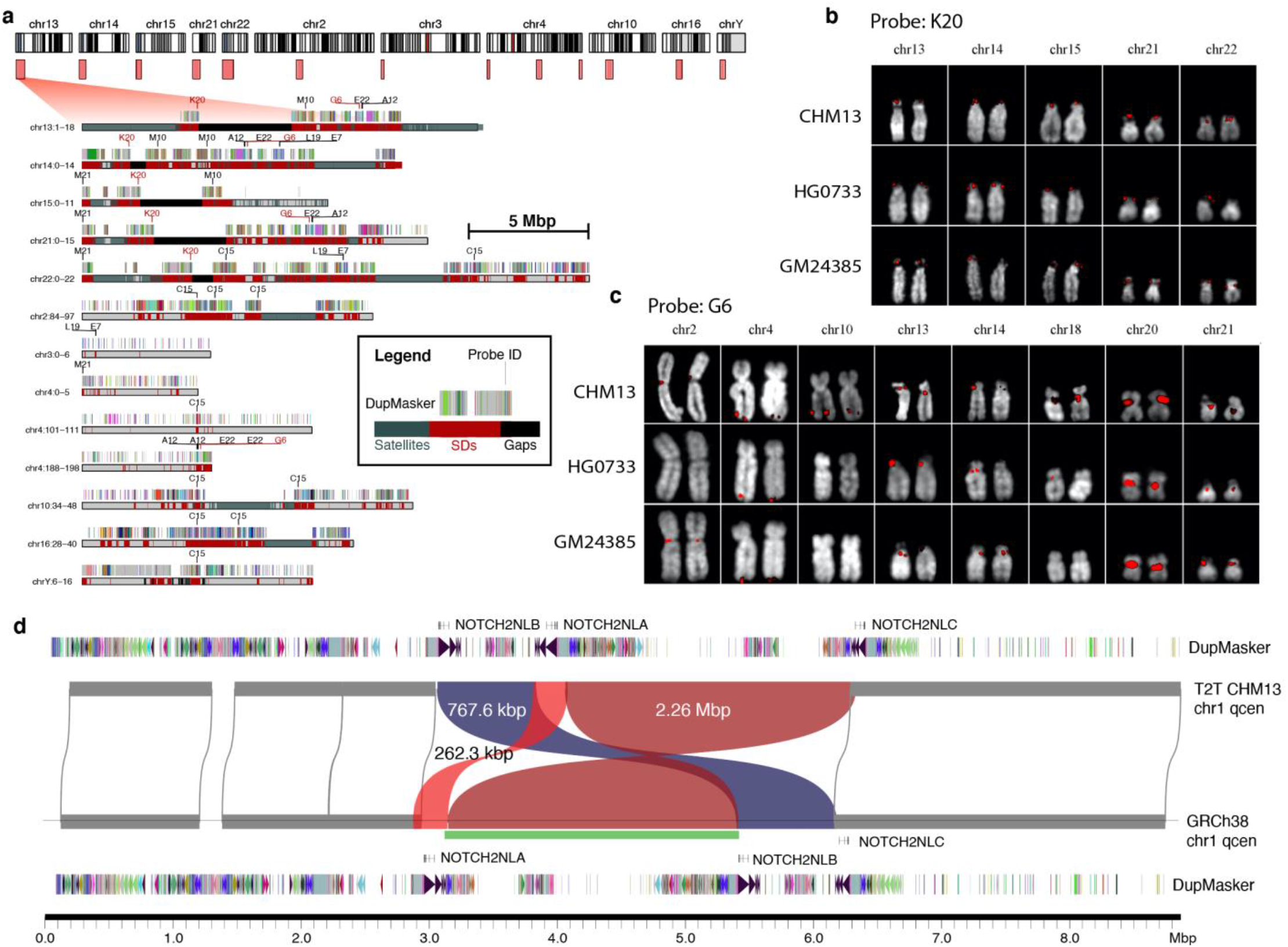
Validation of novel SDs in T2T-CHM13 and heteromorphic variation. **a)** Ideogram (top) depicts large SD regions (light red boxes) present in T2T-CHM13 but absent from the current reference human genome (GRCh38). An expanded view of the duplication (red) and satellite organization (blue-gray) are depicted below showing the location of fosmid FISH probes (e.g., C15) and SD organization compared to ancestral duplicon segments (multi-colored bars) (see inset). **b**,**c)** FISH signals (red) shown on extracted metaphase for two probes and three human cell lines. Probe K20 shows a fixed signal (except for one heterozygous signal), and G6 is heteromorphic among humans (see Table S4, Fig. S4 for complete description for all nine probes). **d)** Inversion polymorphism (green bar) between T2T-CHM13 and GRCh38 in the pericentromeric chromosome 1q region. The inversion (green bar) identified by Strand-seq (*29*) is confirmed in the assembly; however, the sequence-resolved assembly shows a more complex structure including two inversions (red) and one reordered segment (blue) mapping near the *NOTCH2NL* human-specific duplications.

There is an excellent correlation (Pearsons=0.96) between genome-wide copy number variation from the assembly and Illumina read-depth data generated from the same CHM13 source (Methods). Because SDs frequently map to the breakpoints of inversion polymorphisms (*25, 29, 30*), we validated 65 inversions relative to GRCh38 based on Strand-seq analysis of the CHM13 assembly (Fig. S5-S6, Methods). While 32 of these represent known human polymorphisms, 33 are novel compared to six previously analyzed human genomes (*29*). However, by analysis of Strand-seq data from one additional human haplotype (CHM1), we further confirmed 30 of these inversions (i.e., present in CHM1 and CHM13) suggesting that at least 95.4% (62/65) represent true large-scale human inversion polymorphisms (Fig. S5). The inversions associated with SDs (30) are significantly longer than those not associated with SDs (p-value < 0.01, one-sided Wilcoxon rank-sum test) and are all polymorphic among humans (Fig. S6). One striking example is an inversion polymorphism mapping to chromosome 1q21. It is a complex event consisting of two inversions (262.3 kbp, 2.26 Mbp) originally predicted by Sanders and colleagues (*30*) but our sequence analysis shows a relocation of 767.6 kbp of genic sequence (Fig. 2d). The large inversion (chr1:146,350,000-148,610,000) is flanked by the core duplicon, the *NPBF* gene family, and in combination with the other rearrangements changes the order of human-specific genes *NOTCH2NL*A, B and C, which have been implicated in the expansion of the frontal cortex (*8, 31*). As a final test, we sequence resolved this region in eight additional human haplotypes (Methods)—all of which support the T2T-CHM13 configuration with one exception (CHM1), which is consistent with the GRCh38 configuration (Fig. S7).

### Single-nucleotide and copy number variation within SDs

The high quality and single haplotype nature of both the T2T-CHM13 and GRCh38 reference genomes provides us an opportunity to compare the genome-wide pattern of single-nucleotide variation in regions that have been typically excluded from most previous analyses due to their repetitive nature. We aligned GRCh38 to T2T-CHM13 in 5 kbp windows and retained only regions deemed to be “syntenic” based on an unambiguous one-to-one correspondence between both reference genomes (Methods). Most unique regions of the genome (2,693 Mbp) could be compared while only 60% (124 Mbp) of the SDs within T2T-CHM13 had a clear orthologous relationship between the two human references. As expected, the X chromosome and the region corresponding to the major histocompatibility complex (MHC) are the least and most divergent, respectively (Fig. 3a), due to the slower rate of evolution for the female X and the deep coalescence of MHC. Of note, SD sequences are significantly more diverged than unique sequences (p-value < 0.001, one-sided Mann-Whitney U test) (Fig. S8). Comparing only syntenic regions between GRCh38 and T2T-CHM13, we estimate the single-nucleotide variant (SNV) density to be 0.95 SNVs/kbp for unique regions of the genome when compared to SD regions where density rises to 1.47 SNVs/kbp (Table S5). This 50% increase could be due to an increased mutation rate of SDs (e.g., due to the action of interlocus gene conversion), or a deeper average coalescence of duplicated sequences.

**Fig. 3.**
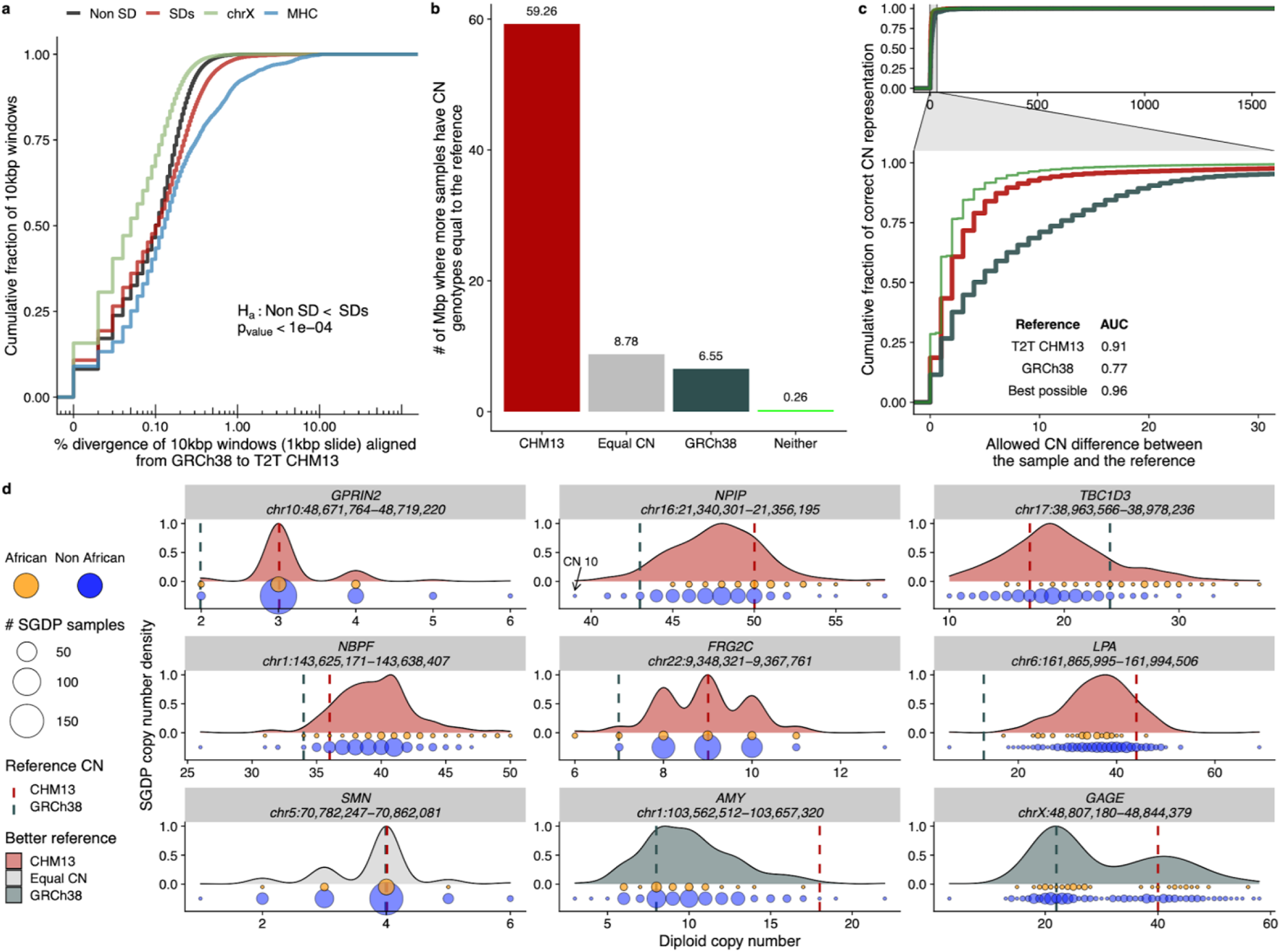
SD single-nucleotide and copy number variation. **a)** Sequence divergence (% in 10 kbp bins) based on syntenic alignments between GRCh38 and T2T-CHM13 for SDs (red), and unique genomic regions (black). SD regions show significantly more divergence when compared to unique sequence (black) and chromosome X (blue) but less than the MHC regions (green). **b)** Copy number (CN) of SD regions that are new or structurally different in T2T-CHM13 compared to GRCh38 based on 268 human genomes from the Simons Genome Diversity Project (SGDP). The histogram shows the number of Mbp where more samples support the CN of the given assembly [T2T-CHM13 (red), GRCh38 (blue), neither (green), or both equally (Equal CN)]. **c)** Empirical cumulative distribution showing how many samples genotype correctly with either GRCh38 or T2T-CHM13 as a function of the allowed difference between sample and reference CN. The inset shows the area under the curve (AUC) calculation for both references allowing a maximum CN difference of 30. The green curve shows an *in silico* reference made using the median CN of the SGDP samples at each site. **d)** Genic copy number variation. Copy number variation in nine gene families are shown (based on SGDP) and distribution is colored according to which reference better reflects the median CN; GRCh38 generally underestimates CN (vertical lines) and Africans (orange) tend to show higher CN than non-Africans (blue); circle size indicates # of samples.

As part of this analysis, we also identified regions that structurally differ or are absent from GRCh38 when compared to the T2T-CHM13 assembly. Based on 1 Mbp LASTZ alignments (Methods), we identified 126 non-syntenic regions for a total of 240 Mbp (N50 length of 12.7 Mbp; Fig. S9). Of these, 33.9% (81.34/240 Mbp) overlap SD regions. Using sequence read-depth (Methods) from 268 human genomes (Simons Genome Diversity Project or SGDP), we compared the copy number of both T2T-CHM13 and GRCh38 (*32*) successfully genotyping 1,292 distinct copy number variable regions (74.85 Mbp). We find that CHM13 approximates the median (+/-2 s.d.) human copy number (CN) from SGDP for 94% of bases (70.6 Mbp) in contrast to GRCh38 where 57% of bases (42.8 Mbp) meet this metric (Fig. S10). In particular, we find that human copy number is nine times (59.26/6.55 Mbp) more likely to match the CHM13 copy number rather than GRCh38 (Fig. 3b). Thus, CHM13 is a much better predictor (AUC 0.91) than GRCh38 (AUC 0.77) of human copy number variation and better approximates an *in silico* human reference constructed using the median CN of the SGDP samples at every site (AUC 0.96, Fig. 3c). GRCh38 tends to underestimate normal human CN (by on average 9.2 copies or a median of 3.0 copies).

We identify 119 protein-encoding genes (65 for GRCh38) where CHM13 copy number better represents the true human copy number state (Table S6). These include both biomedically important genes relevant to disease risk *(LPA, MUC3A, FCGR2*) (*33*–*41*) as well as gene families that have been implicated in the expansion of the human brain during human evolution (*TBC1D3, NPIP, NPBF*) (Fig. 3d, Table S6) (*7, 42*–*44*). In T2T-CHM13, for example, there are additional copies of *NPIP, NPBF*, and *GOLGA* that are absent from GRCh38—each of these has been described as core duplicons responsible for the expansion of interspersed duplications in the human genome (*24*) as well as the emergence of human-specific gene families. Interestingly, African genomes tend to have overall a higher copy number status when compared to non-African genomes. In particular, *TBC1D3* shows ∼7 fewer copies in non-Africans when compared to Africans (p-value < 1e-12). These findings suggest that higher copy number is likely ancestral (Table S7) and CHM13, once again, better captures that diversity. Despite its primarily European origin, our results show that the more complete genome assembly serves as a better reference than GRCh38 for copy number variation irrespective of population group (Fig. S22).

### Structural variation and massive evolutionary changes in the human lineage

Advances in long-read genome assembly (*45, 46*) enable sequence resolution of complex structural variation associated with SDs at the haplotype level (*47*). We generated or used existing high-fidelity (HiFi) sequence data from 12 human and 6 nonhuman primate genomes to understand both the structural diversity and evolution of specific SD regions. Comparing the chimpanzee genome (Methods) and the T2T-CHM13 assemblies, we specifically searched for gene-rich, large-scale genetic differences (>50 kbp in length) and selected 10 loci for a more detailed evolutionary analysis, including regions of biomedical importance and regions associated with the expansion of the human frontal cortex (Tables S8-S10; Fig. S11). Of 10 targeted loci, assemblies of additional haplotypes recapitulated the structural organization of T2T-CHM13 for eight of the 10 loci whereas evidence for the structural organization of GRCh38 was only found in five of the 10 loci (Methods). Overall, 73% of human haplotype assemblies were successfully reconstructed (Table S8); however, the loci varied depending on the size and complexity of the locus. For example, in the case of the 8.9 Mbp region corresponding to *NOTCH2NL* and *SRGAP2B/2D*, we recovered only 37.5% of human haplotypes (Table S8, Fig. S7). Similarly, we resolved only six haplotypes (from a potential of 24 haplotypes) for the 3.4 Mbp region harboring the *SMN1 and SMN2* loci (Fig. S12).

Among haplotypes that could be resolved, we find a high degree of structural heterozygosity among human genomes (67%, Methods) with 249 kbp differing on average when compared to T2T-CHM13 (Table S9). In some cases, the structural changes are simple, such as ∼12 kbp insertion or deletion of *CYPD26*, which contributes to differential drug metabolism activity as well other human disease susceptibilities (*48*–*54*) (Fig. S13). In other cases, the patterns of structural variation are complex involving hundreds of kilobase pairs of inserted or deleted gene-rich sequence along with large-scale inversion events that alter gene order for specific human haplotypes (see *ARGHAP11A/B*; Fig. S14 and *NOTCH2NLA/B;* Fig. S7). The spinal muscular atrophy (SMA) locus containing *SMN1* and *SMN2—*one of the most difficult regions to finish as part of the human genome project on chromosome 5 (*55*)—shows a unique structure for each of the six assembled haplotypes that we resolved (plus GRCh38). Some haplotypes not only show increases in *SMN2* copy number (Fig. S12), a known genetic modifier of SMA (*56*), but also potential functional differences in the organization and composition of *SMN2*. Since *SMN2* serves as a target for small-molecule drug therapy improving splice-site efficiency compensating for the loss *SMN1* in SMA patients (*57*), this level of sequence resolution is of practical utility for disease risk assessment and treatment of patients.

Of particular interest is the *TBC1D3* gene family (*42*) (Fig. 4, Fig. S15-S16) whose protein products modulate epidermal growth factor receptor signaling and trafficking (*58*) and whose duplication in humans has been associated with expansion of the human prefrontal cortex as evidenced by mouse transgenic experiments (*7*). A comparison to chimpanzee (Fig. 4a) shows two massive genomic expansions in the human lineage (323.0 and 124.4 kbp). Both the high sequence identity (99.6%) and sequence read-depth comparisons of *TBC1D3* copy number are consistent with expansion occurring in the human lineage after divergence from chimpanzee (Fig. 4b). We extended this analysis to other nonhuman primates by generating HiFi assemblies for bonobo, gorilla, orangutan, and macaque. We identified *TBC1D3* homologues in each species and constructed a maximum likelihood phylogeny based on intronic or noncoding sequence flanking the gene (Fig. 4c). The analysis reveals recurrent and independent expansions of *TBC1D3* in the orangutan, gorilla, and macaque species at different time points during primate evolution with the most recent occurring 2 and 2.6 million years ago, near the emergence of the *Homo* genus (*59*).

**Fig. 4.**
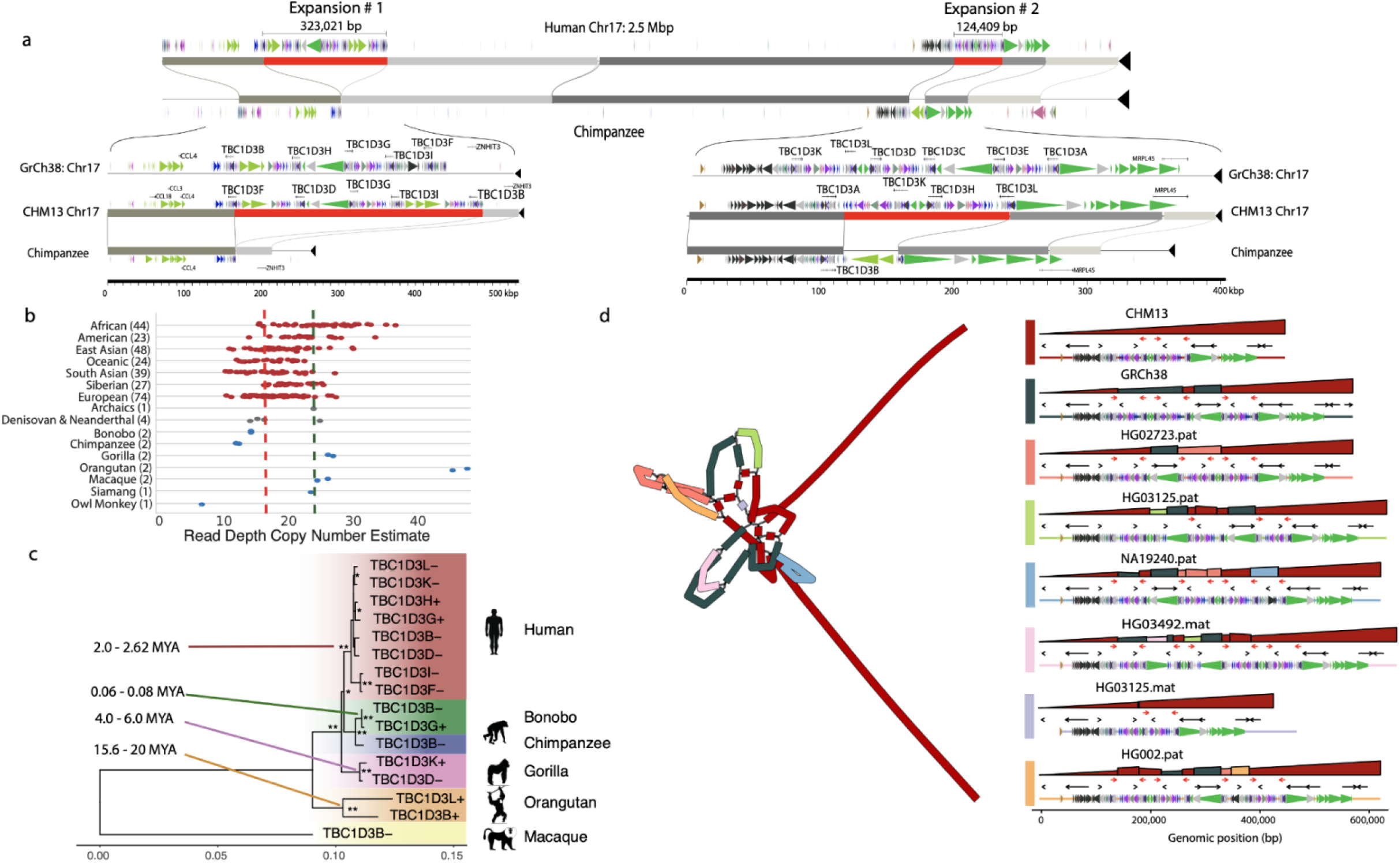
Human-specific expansion of TBC1D3 compared to nonhuman primates. **a)** Regions of homology between human T2T-CHM13’s Chromosome 17 (top) and a HiFi assembly of the chimpanzee genome (bottom). Red blocks represent regions of human-specific expansion, including *TBC1D3* duplications. Colored arrows above and below the homologous sequence represent unique ancestral units (duplicons) identified by DupMasker. Inset plots for both expansion sites are included below with the gene models identified by Liftoff (*89*). **b)** CN (diploid) estimates from an Illumina read-depth analysis of SGDP, ancient hominids, and nonhuman primates for a *TBC1D3* paralog. CN estimates include pseudogenes (5) not included in the phylogeny, explaining the higher counts observed. The T2T-CHM13 CN and GRCh38 CN are represented by the red and blue lines, respectively. **c)** Phylogeny of *TBC1D3* copies at these two expansion sites as well as nonhuman primate copies. Single asterisks at nodes indicate bootstrap values greater than or equal to 70%, while double asterisks indicate 100%. The data illustrate a human-specific expansion, as well as several independent expansions in the macaque, gorilla, and orangutan. Using macaque sequence as an outgroup, we estimate the human-specific expansion to be ∼2.3 million years ago (MYA). **d)** Variation in human haplotypes across the first *TBC1D3* expansion site: a graph representation (rGFA, left) of the locus where colors indicate the source genome for the sequence, and on the right the path for each haplotype-resolved assembly through the graph. The top row for each haplotype composed of large polygons represents an alignment comparing the haplotype-resolved sequence (horizontal) against the graph (vertical), and color represents the source haplotype for the vertical sequence. For example, a single large red triangle indicates there is a one-to-one alignment between CHM13 and the haplotype. Structural variants can be identified from discontinuities in height (deletion), changes between colors (insertion), or changes in the direction of the polygon (inversion). Below is shown the gene of interest (red arrow) and other genic content in the region (black arrow). Colored bars show ancestral duplication segments (duplicons) that compose the larger duplication blocks.

Complete sequencing of human *TBC1D3* haplotypes reveals remarkable structural diversity (Fig. 4d) with *TBC1D3* copy number ranging from three to fourteen *TBC1D3* copies at expansion site #1, and two to nine copies at expansion site #2. In total, approximately one-third of human expansion site #2 shows large-scale structural variation and we identify >1.8 Mbp of duplicated sequence and >650 kbp of inverted sequence across the 18 haplotypes (including GRCh38). We estimate the heterozygosity of this locus to be over 77.8% (14/18 haplotypes are structurally distinct) (Fig. S16). Similarly, *TBC1D3* expansion site #1 is structurally heterozygous with 63.6% (14/22) of the haplotypes displaying unique structures corresponding to copy number differences in the *TBC1D3* gene family (Fig. S15). Using orthogonal Oxford Nanopore Technologies (ONT) ultra-long-read sequencing, we validated these complex patterns of structural variation in a subset of the samples investigated here (Methods, Fig. S17-S18). To better represent the structural genetic variation at this locus, we used a graph-based representation (*60*), which identified two *TBC1D3* genes as common among all human haplotypes examined thus far (*TBC1D3B* at site #1 and *TBC1D3A* at site #2).

### New gene models and variable duplicate genes

We identified 182 candidate new or non-syntenic genes (Supplemental note 2) in the T2T genome assembly (compared to GRCh38) with open reading frames and multiple exons (Table S11). Of these 91% (166) corresponded to SD gene families (Fig. 5a). Many of these represent expanded tandem duplications (e.g., *GAGE* gene family members on the X chromosome) or large interspersed duplications (e.g., beta-defensin locus) adding additional copies of nearly identical genes to the human genome (Fig. 5a). We searched for evidence that these copy number polymorphic or structurally variant regions were transcribed by aligning long-read transcript sequencing data and searching for perfect matches (Methods). We constructed a database of 44.2 million full-length cDNA transcripts derived from 31 human tissue samples and compared them to both the GRCh38 and T2T-CHM13 human genome references. For those 182 novel protein-coding genes where an unambiguous assignment could be made, 36% (65/182, >20 Iso-Seq reads) were confirmed to be expressed with 23 showing the majority of reads mapping better to T2T-CHM13 when compared to GRCh38 (Fig. 5b). Overall, across the entire genome, 12% of full-length transcripts exhibit at least 0.2% higher alignment identity when mapped against CHM13, while 8% align better to GRCh38. These results are consistent with the notion that the T2T-CHM13 is more complete, but that both assemblies are in some cases capturing different structurally variant haplotypes associated with genes. In addition to entirely new genes, we identify several gene models that are complete for the first time—many of which encode proteins with large tandem repeat domains (ZNF, LPA, Mucin; Fig. 5c). Among these is the complete gene structure of the Kringle IV domain of the lipoprotein A gene. Reduced copies of this domain are among the strongest genetic associations with cardiovascular disease, especially among African Americans (*33*–*36, 61*). Sequencing of multiple human haplotypes not only identified length variation but also other forms of rare coding variants potentially relevant to disease risk (Fig. 5d).

**Fig. 5.**
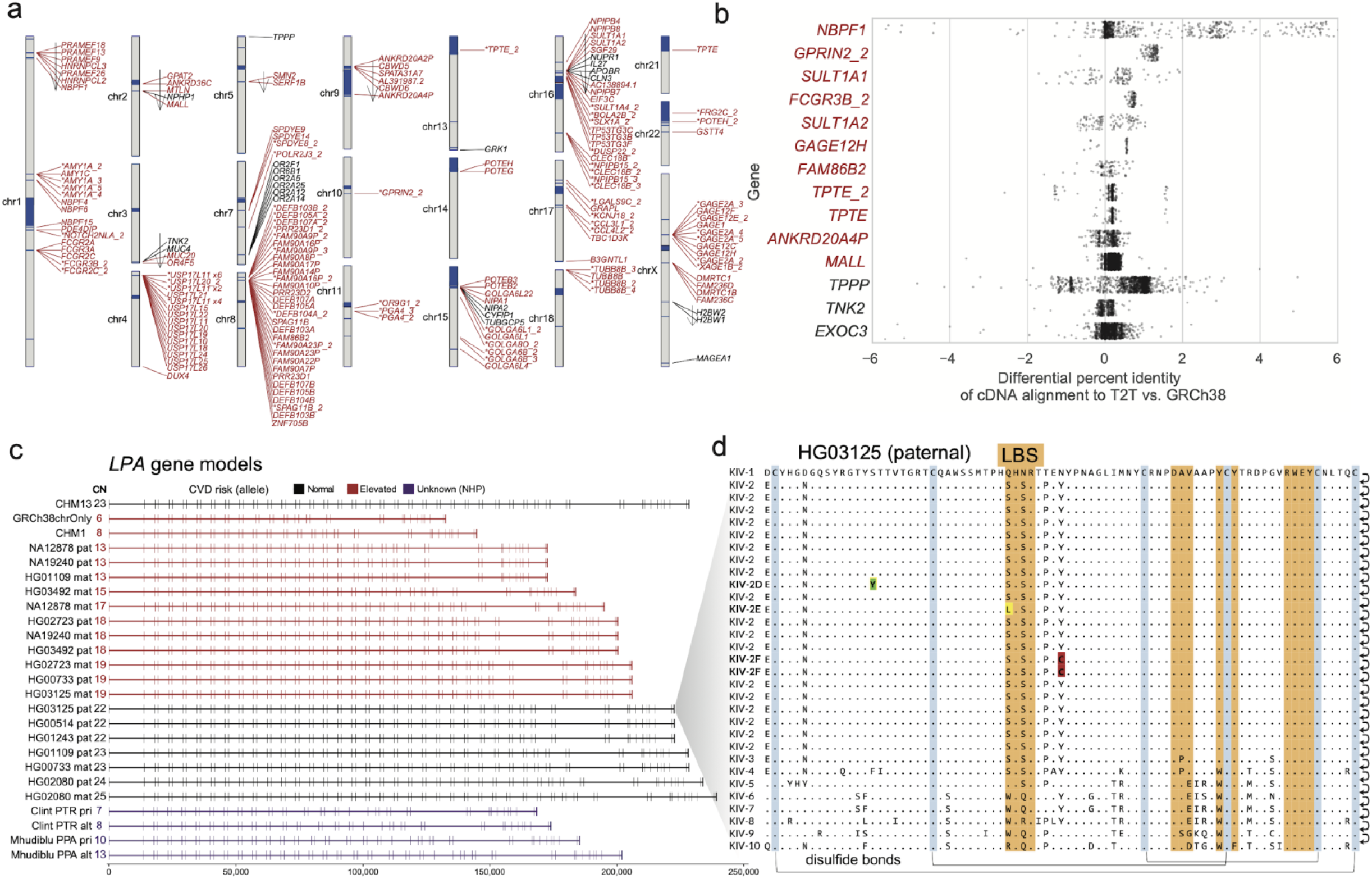
Genic variation in new SD regions of T2T-CHM13. **a)** Ideogram showing the new or non-syntenic gene models (open reading frames [ORFs] with >200 bp of coding sequence and multiple exons) in the T2T-CHM13 assembly as predicted by Liftoff. New genes mapping to SDs (red) are indicated with an asterisk if predicted to be an expansion in the gene family relative to GRCh38 (Methods). Arrows indicate inverted regions. Most unique genes mapping to non-syntenic regions (black) are the result of an inversion (arrow). **b)** Percent improvement in mapping of CHM13 Iso-Seq reads in candidate duplicated genes (red) mapping to non-syntenic regions of the T2T-CHM13 assembly. Positive values identify Iso-Seq reads aligning better to T2T-CHM13 than GRCh38. **c)** Gene models of *LPA* with ORF generated from haplotype-resolved HiFi assemblies. The double-exon repeat in these gene models encode for the Kringle IV subtype 2 domain of the LPA protein. Highlighted in red are haplotypes with reduced Kringle IV subtype 2 repeats predicted to increase risk of cardiovascular disease. **d)** Amino acid variation in the Kringle IV subtype 2 repeat in the paternal haplotype of HG01325 identifies a previously unknown set of amino acid substitutions including rare variants: Ser42Leu in the active site, Ser24Tyr and Tyr49Cys.

### SD methylation and transcription

Since methylation is an important consideration in regulating gene transcription, we took advantage of the signal inherent in ultra-long-read ONT data (*62*–*64*) to investigate the CpG methylation status of SD genes within the CHM13 genome (Methods). Using hierarchical clustering, we find that SD blocks are generally either methylated or unmethylated as an entire block; (Fig. S19, Fig. 6a). Specifically, we find that 452 SD blocks flanked (127.7 Mbp) by unique sequences are hypermethylated in contrast to 222 hypomethylated SD blocks (52.1 Mbp). Methylation status does not appear to be driven by genomic location, e.g., proximity to the centromeres, acrocentric short arms, or telomeres (Fig. 6a). Using full-length transcript data from CHM13, we compared methylation and transcription status of duplicated genes (Methods). If we stratify genes by their number of full-length transcripts, we observe distinct methylation patterns for transcribed and non-transcribed SD genes (Fig. 6b). For highly transcribed SD genes and unique genes (genes without at least one exon overlapping with SD sequence), the gene body and flanking sequence are generally hypermethylated with a dramatic dip near the transcription start site (TSS)/promoter (*65*). In contrast, non-transcribed genes show moderate to low methylation across the gene body and flanking sequence. Restricting the analysis to genes mapping within SDs, we find that transcriptionally silenced duplicate genes are more likely (10,000 permutations, p=0.0018) to map to hypomethylated regions of SD sequence (Fig. 6a) when compared to transcribed duplicate genes. Additionally, in untranscribed SD genes we observe a statistically significant (one-sided Mann-Whitney) increase in TSS methylation (6.6% increase) when compared to unique genes where the TSS is more likely to be depleted for methylation (8.2% decrease).

**Fig. 6.**
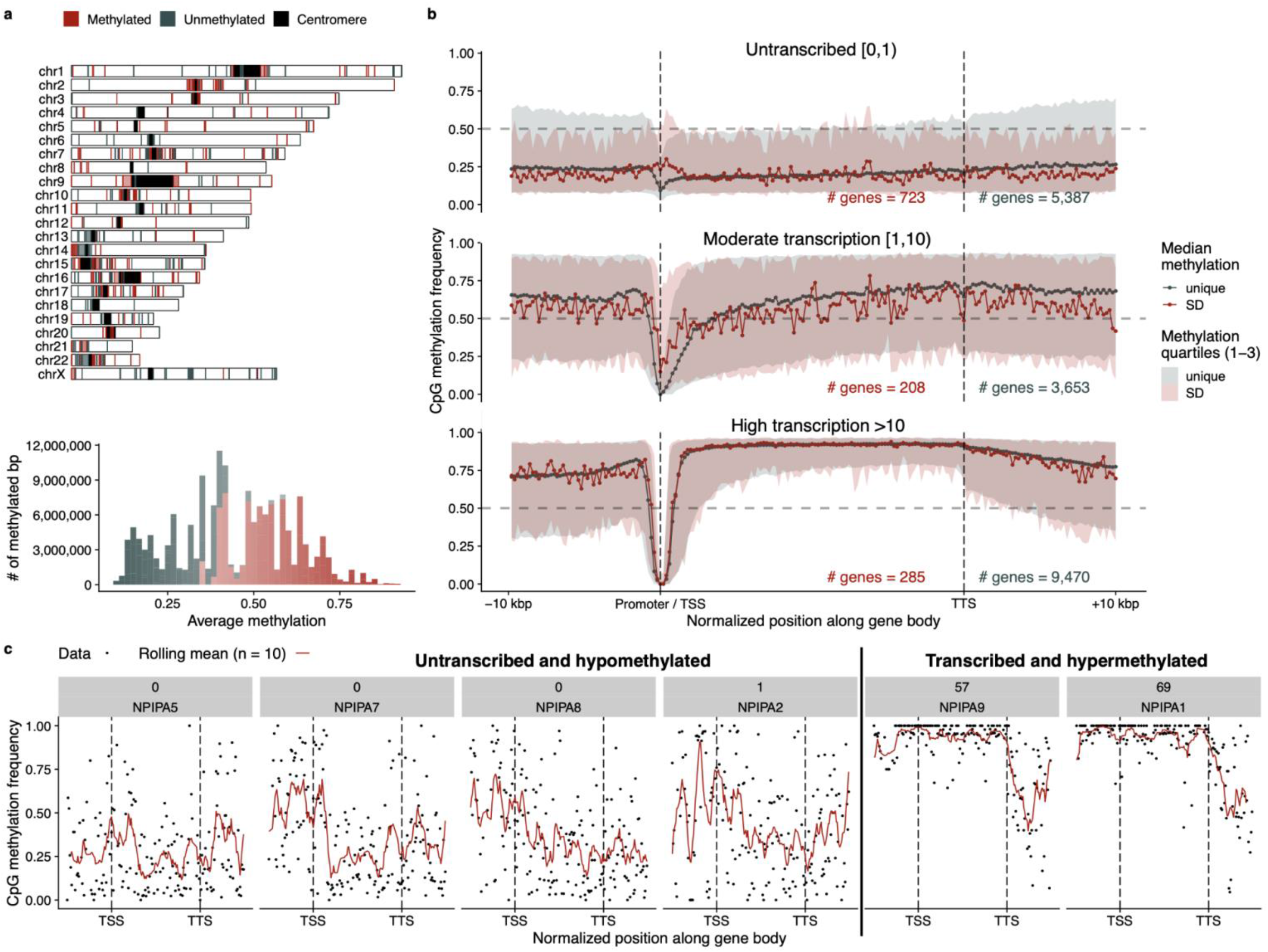
SD methylation and gene transcription. **a)** Methylated (red) or unmethylated (blue-gray) SD blocks in the CHM13 genome based on processing ONT data. The histogram shows the distribution of average methylation across these regions. **b)** Median methylation signal of SD (red) and unique (blue-gray) genes stratified by their Iso-Seq expression levels in CHM13. The filled intervals represent the 25 and 75 quartiles of the observed data. Vertical lines indicate the position of the transcription start site (TSS) and the transcription termination site (TTS). **c)** Methylation signal across the recently duplicated *NPIPA* gene family in CHM13, showing increased methylation in transcriptionally active copies. Black points are individual methylation calls, and the red line is a rolling mean across 10 methylation sites. The labels in gray show the number of CHM13 Iso-Seq transcripts and the gene name.

One important consideration in this analysis is the presence of a CpG island within 1500 bp of the promoter (*66*). In our analysis of CHM13, for example, unexpressed unique genes have a low CpG count, consistent with a lack of CpG islands (Fig. S20). If we repeat the same analysis on SD genes, we find that the unexpressed SD genes exist with and without CpG islands (Fig. S20). In total, these observations suggest a process of epigenetic silencing for a subset of duplicate genes through general demethylation of the gene body but hypermethylation of promoter regions. Based on these observed signatures, we investigated whether it might be possible to predict actively transcribed duplicate gene copies from these epigenetic features. We investigated a recently duplicated hominid gene family (*NPIPA*) (*67*) where sufficient paralogous sequence differences exist to unambiguously assign full-length transcripts to specific loci. While promoter/TSS signatures are less evident at the individual gene level, the gene body methylation signal appears diagnostic (Fig. 6c). *NPIPA1* and *NPIPA9*, for example, are the most transcriptionally active and show demonstrably distinct methylation patterns providing an epigenetic signature to distinguish transcriptionally active loci associated with high-identity gene families that are otherwise largely indistinguishable. We show this trend also holds for other high-copy number gene families (Fig. S21).

## DISCUSSION

This work provides the first comprehensive view of the organization of SDs in the human genome. The new reference adds a chromosome’s worth (81 Mbp) of new SDs increasing the human genome average from 5.4 to 7.0% nearly doubling the number of SD pairwise relationships (24 vs. 41 thousand) and, as a result, predicts new regions of genomic instability due to their potential to drive unequal crossing-over events during meiosis. By every metric, T2T-CHM13 is a better representation of the structure of the human genome than GRCh38. This includes the first sequence-based organization of the short arms of chromosomes 13, 14, 15, 20 and 21 where we find that SDs account for more sequence (34.6 Mbp) than either heterochromatic satellite (26.7 Mbp) or rDNA (10 Mbp). Acrocentric SDs are almost twice as large when compared to non-acrocentric regions likely due to ectopic exchange events occurring among the short arms, which associate more frequently during the formation of the nucleolus (*68*). Interestingly, nearly half of the acrocentric SDs involve duplications with non-acrocentric pericentromeric regions of chromosomes 1, 3, 4, 7, 9, 16, and 20. These duplicated islands of euchromatic-like sequences within acrocentric DNA are much more extensive than previously thought but have been shown to be transcriptionally active (*69*). While the underlying mechanism for their formation is unknown, it is noteworthy that three of the non-acrocentric regions have large secondary constriction sites (chromosomes 1q, 9q, and 16q) composed almost entirely of heterochromatic satellites (HSAT2 & 3) (Fig. S2). These particular SD blocks, thus, are bracketed by large tracts of heterochromatic satellites and such configurations may make them particularly prone to double-strand breakage events (*70*) promoting such large interchromosomal duplications (Fig. S3) between acrocentric and non-acrocentric chromosomes.

The new T2T-CHM13 reference along with resources from other human genomes provides a baseline for investigating more complex forms of human genetic variation. For example, this truly complete reference sequence facilitates the design of sequence-anchored probes to systematically discover and characterize SD heteromorphic variation where chromosome organization differs among individuals (Fig. 2). Such chromosomal heteromorphisms have been traditionally investigated cytogenetically and are thought to be clinically benign (*26*–*28*); however, more recent work indicates that these large-scale variants associate with infertility by increasing sperm aneuploidy, decreasing rates of embryonic cleavage (IVF), and increasing miscarriages (*71*–*78*). Distinguishing between fixed and heteromorphic acrocentric SDs will facilitate such research as well as the characterization of breakpoints associated with Robertstonian translocations—the most common form of human translocation (*79*).

At a finer-grained level, the new reference and the use of long reads from other human genomes provides access to other complex forms of variation involving duplicated gene families. Short-read copy number variation analyses and single-nucleotide polymorphism microarray have long predicted that SDs are enriched 10-fold for copy number variation but the structural differences underlying these regions as well as their functional consequences have remained elusive (*10, 80*). We reveal unprecedented levels of human genetic variation in genes important for neurodevelopment (*TBC1D3*) and human disease (*LPA, SMN*). Even between just two genomes (GRCh38 and CHM13), we find that 37% (81 Mbp) of SD bases are new or structurally variable and this predicts 182 copy number variable genes between two human haplotypes (Table S6). In cases such as *TBC1D3*, we find that most human haplotypes vary (64-78%). Different humans carry radically different complements and arrangements of the *TBC1D3* gene family. The potential ramifications of this dramatic expansion in humans versus chimpanzees and of such high structural heterozygosity among humans are intriguing given the gene’s purported role in expansion of the frontal cortex (*7*). Similarly, we were able to reconstruct the complete structure of the *LPA* gene model in multiple human genotypes. While this is only a single gene, variability in the tandemly repeated 5.2 kbp protein-encoding Kringle IV domain underlies one of the most significant genetic risk factors for cardiovascular disease. Sequence resolution of the structural variation, as well as underlying amino-acid differences, allow us to predict novel risk alleles for disease (Fig. 5). Sequence-resolved structural variation improves genotyping and tests of selection (*47, 81, 82*) providing a path forward for understanding the disease and evolutionary implications of these complex forms of genetic variation.

Finally, and perhaps most importantly, the new reference coupled with other long-read datasets enables genome-wide functional characterization of recently duplicated genes. Both gene annotation and large-scale efforts to characterize the regulatory landscape of the human genome have typically excluded repetitive regions, including the 859 human genes mapping to high-identity SDs (*83, 84*). This is because the underlying short-read sequencing limits conventional RNA-seq or Chip-seq data from being assigned unambiguously to specific duplicated genes. In this study, we generated long-read full-length transcript data (Iso-Seq) with long-read methylation data from ONT sequencing of the same genome to simultaneously investigate epigenetic and transcriptional data against a fully assembled reference genome. The long-read data from the same haploid source facilitated the unambiguous assignment of these functional readouts allowing us to correlate methylation and transcript abundance. Our initial analyses suggest that a large fraction of duplicate genes are in fact epigenetically silenced (characterized by hypermethylation of the promoter and hypomethylation of the gene body) and that this epigenetic mark may be used to predict actively transcribed loci even when genes are virtually identical (Fig. 6 and S21). While more human genomes and diverse tissues need to be interrogated to assess the significance of this observation, it is clear that phased genome assemblies (*47*) with long-read functional readouts such as methylation (*64*), transcription, or Fiber-seq (*85, 86*) provide a powerful approach to understanding the regulatory landscape of duplicated and copy number polymorphic genes in the human genome.

There are several remaining challenges. First, not all human haplotypes corresponding to specific duplicated regions could be fully sequence resolved using automated assembly of long-read HiFi sequencing technology. The majority of the 250 unresolved regions of phased human genomes generated solely with HiFi long reads correspond to some of the largest and most variable duplicated regions of the human genome (*47*). For example, only 25% of *SMN1*/*SMN2* haplotypes were fully resolved by HiFi assembly and unresolved loci are predicted to carry some of the most complex structural variation patterns. In comparison, the T2T-CHM13 assembly used both accurate HiFi and ultra-long ONT data, and future assembly methods that combine these technologies will likely be critical for diploid T2T assembly and the complete characterization of SD haplotypes (*87, 88*). Another important challenge going forward will be how to accurately represent these more complex forms of human genetic variation, including functional annotation, where linear representations may be insufficient. While a more complex pangenome reference graph could overcome these limitations, it is unclear how this will be achieved in practice or how it will be adopted by the genomics and clinical communities. This highlights the importance of not only the construction of a pangenome reference but the necessary tools that will distinguish paralogous and orthologous sequences within duplications to allow for comparison between haplotypes with different SD architectures. The work currently underway by the Human Pangenome Reference Consortium (HPRC), Human Genome Structural Variation Consortium (HGSVC), and Telomere-to-Telomere (T2T) Consortium will be key to developing these methods and completing our understanding of SDs and their role in human genetic variation.

## METHODS

### Estimating the number of rDNA copies in the assembly

To estimate the CN in the assembly of the rDNA repeats we aligned KY962518.fasta (https://www.ncbi.nlm.nih.gov/nuccore/KY962518) to the whole-genome assembly using minimap2 with the following settings and counted the number of alignments:

~~~
minimap2 -ax asm20 -N 100 -p 0.5 --secondary=yes --eqx -r 500000
chm13.draft_v1.0.fasta KY962518.fasta | samtools view -c
~~~

### SD annotation

To annotate SDs we identified homologous segments using SEDEF [v1.1-31-g68de243, (*19*)] on a masked version the T2T-CHM13 v1.0 assembly that included chrY from GRCh38. SDs were filtered to contain at most 70% satellite sequence as determined by RepeatMasker. Additionally, SDs had to be at least 90% identical by gap-compressed identity, 50% identical including indels (blast identity), and at least 1 kbp of aligned sequence or else they were filtered into a set of smaller and lower identity duplications. Pericentromeric and telomeric SDs were defined as being within 500 kbp and 5 Mbp of the telomere and centromere, respectively. The full pipeline for these masking steps is provided for convenience at https://github.com/mrvollger/assembly_workflows/ under workflows/sedef.smk. The same workflow was applied to the chromosome-level scaffolds of GRCh38 for all SD comparisons made in the paper.

### Repeatmasking

Common repeats were masked with RepeatMasker v4.1 (*90*) and Tandem Repeats Finder (TRF) (*91*). The full pipeline for these masking steps is provided for convenience at https://github.com/mrvollger/assembly_workflows/ under workflows/mask.smk. In brief, RepeatMasker was run with the following settings:

~~~
 RepeatMasker \
  -s \
  -xsmall \
  -e ncbi \
  -species human \
  -dir $(dirname {input.fasta}) \
  -pa {threads} \
  {input.fasta}
~~~

And TRF was run with

~~~
 trf {input.fasta} 2 7 7 80 10 50 15 -l 25 -h -ngs > {output.dat}
~~~

### Defining syntenic regions between T2T-CHM13 and GRCh38

The T2T-CHM13 to GRCh38 synteny track was constructed using the [Cactus HAL file](https://github.com/ComparativeGenomicsToolkit/cactus#using-the-output) (available at the following link: http://t2t.gi.ucsc.edu/chm13/hub/t2t-chm13-v1.0/cactus/t2t-chm13-v1.0.aln1.hal) with 1 Mbp resolution and a maximum anchor distance of 50 kbp. We used the tool halSynteny to construct syntenic blocks from the Cactus alignments—the methods of which are described in detail in Krasheninnikova et al. (*92*). This track is available at the following link: http://t2t.gi.ucsc.edu/chm13/hub/t2t-chm13-v1.0/synteny/synteny.1mb.bigPsl. To define the new and variable regions of T2T-CHM13, we inverted the 1 Mbp synteny track retaining all regions without an alignment to GRCh38.

### Calculating the number of SD alignments in 5 Mbp windows

We first offset the SD coordinates in GRCh38 such that the largest gaps (acrocentric short arms, centromeres, and HSAT arrays) matched the length of the assembled sequence in T2T-CHM13. We then normalized the GRCh38 coordinates so that the length of the chromosomes in GRCh38 were equal to those in T2T-CHM13. After this we took 5 Mbp non-overlapping windows from T2T-CHM13 and the normalized GRCh38 and calculated the difference in the number of SDs within each window (Table S3).

### WSSD detection and genotyping

As an orthogonal method to estimate copy number of SDs, we applied the whole-genome shotgun sequence detection (WSSD) pipeline, which uses sequence read-depth as a proxy (*14*). Short-read sequence data were processed into 36 bp non-overlapping fragments and mapped to a masked T2T-CHM13 reference using mrsFAST (*93*) with a maximum of two substitution mismatches not allowing for indels. Masking was determined by TRF and RepeatMasker. Read-depth across the genome was corrected for GC-bias and copy number was determined using linear regression on read-depth versus known fixed copy number control regions. Finally, integer genotypes were estimated by using the predicted mean and variance of the Gaussian distributions underlying different copy numbers to create a series of models to represent the likely distribution of read depths underlying a region of specific copy number.

For defining genotyping intervals, we applied the changepoint package in R (*94*) to identify regions where the CHM13 WSSD CN estimate was consistent. Specifically, we used a log-transformed continuous CN estimate from WSSD for sliding windows across the assembly and then applied binary segmentation to identify regions where the CN remained the same. We used the following R command:

~~~
cpt.mean(Log_cn, method = “BinSeg”, Q=Q)
~~~

Where `Log_cn` is a vector of log-scaled CN estimates and `Q` is the number of independent 50 kbp windows within each chromosome.

To validate the CN of assemblies we fragmented the assemblies in 36 bp windows with a 1 bp slide and used it as input read data for our WSSD CN pipeline. Then every CN estimate within an SD space was compared between the Illumina estimate and assembly estimate and a Pearson’s correlation was calculated.

### Gene annotations with Liftoff

Gene annotations on T2T-CHM13 were made using Liftoff (*89*) and then processed with gffread (*95*) to filter for only transcripts with open reading frames. The full pipeline for gene annotation is provided for convenience at https://github.com/mrvollger/assembly_workflows/ under workflows/liftoff.smk. In brief, Liftoff was called with the following command:

~~~
 liftoff -dir {output.temp} \
  -f <(echo “locus”) \
  -flank 0.1 \
  -sc 0.85 -copies -p {threads} \
  -g {input.gff} -o {output.gff} -u {output.unmapped} \
  {input.t} {input.r}
~~~

Using as input the GENCODE Genes track v34 annotation gff3 available at ftp://ftp.ebi.ac.uk/pub/databases/gencode/Gencode_human/release_34/gencode.v34.annotation.gff3.gz and GRCh38 FASTA available at https://ftp.ncbi.nlm.nih.gov/genomes/all/GCF/000/001/405/GCF_000001405.39_GRCh38.p13/GRCh38_major_release_seqs_for_alignment_pipelines/GCA_000001405.15_GRCh38_no_alt_analysis_set.fna.gz.

### Counting the number of high-identity SD genes

We counted all protein-encoding genes with at least one exon mapping fully within a >95% identical SD and had the additional condition that at least 50% of the full-length gene maps to SD space without the identity limitation.

### Cell culture

CHM13 and CHM1 cells were cultured in complete AmnioMax C-100 Basal Medium (Thermo Fisher Scientific, 17001082) supplemented with 15% AmnioMax C-100 Supplement (Thermo Fisher Scientific, 12556015) and 1% penicillin-streptomycin (Thermo Fisher Scientific, 15140122). GM24385, GM19240, HG00514 and HG00733 cells were cultured in RPMI 1640 with L-glutamine medium (Thermo Fisher Scientific, 11875093) supplemented with 15% FBS (Thermo Fisher Scientific, 16000-044) and 1% penicillin-streptomycin (Thermo Fisher Scientific, 15140122). All cells were cultured in a humidity-controlled environment at 37°C with 5% CO_2_.

### FISH characterization and validation

Fosmid probes for FISH experiments were selected by mapping fosmid end sequences from the ABC10 (NA19240 Yoruban) library (*25*) to the T2T-CHM13 reference using blast (*96*). Human fosmid clones were used as probes in one- or two-color FISH experiments and hybridized on metaphases obtained from CHM13, CHM1, GM24385, GM19240, HG00514, and HG00733 lymphoblastoid cell lines. FISH experiments were essentially performed as previously described (*97*). Slides were imaged on an inverted fluorescence microscope (Leica DMI6000) equipped with a charge-coupled device camera (Leica DFC365 FX). Mapping was performed following comparison to the conventional classical cytogenetics G-banding (*98*).

### Assembly of additional humans and nonhuman primates

All assemblies with the exception of T2T-CHM13 and GRCh38 were assembled with Hifiasm v0.12 using default parameters. The human samples with the exception of CHM1 were assembled using parental short-read data for phasing. All nonhuman primates and CHM1 were assembled without parental phasing information since none exists.

### ONT validation

To validate structural variant configurations predicted by HiFi sequence and assembly, we aligned ultra-long ONT data from two samples (HG002, HG00733) and assessed the uniformity of coverage over the *TBC1D3* assemblies for these four haplotypes. We find no obvious sign of collapsed duplications (read coverage abnormalities) or misjoins in the assemblies (every 25 kbp segment with 1 kbp slide is spanned by four or more reads) in the ultra-long ONT data (Fig. S18-S19).

### TBC1D3 phylogenetic tree construction

Orthologous sequences for the two human *TBC1D3* expansion sites were identified in T2T-CHM13 using minimap2 (*99*) and gene models were annotated using Liftoff (*89*). *TBC1D3* transcripts with open reading frames were identified using gffread (*95*). Exons were masked and removed using BEDTools maskfasta and getfasta functions (*100*) in order to construct neutrally evolving phylogenetic trees. With exon-free paralogs of both CHM13 and nonhuman primates, a multiple sequence alignment (MSA) was generated using MAFFT (*101*). To produce the most confident MSA, an iterative refinement algorithm described was used with the option for iterating 1000 times (*102, 103*).

~~~
 mafft --reorder --maxiterate 1000 --thread 16 {input.fasta} >
 {output.MSA.fasta}
~~~

The MSA was subsequently used to generate a maximum likelihood phylogeny, using RAxML (*104*). For this phylogeny, the rapid bootstrapping analysis was utilized to identify the best maximum likelihood tree, a gamma model was used to model rate heterogeneity, and macaque *TBC1D3* sequences were used as outgroup sequences.

~~~
 raxmlHPC-PTHREADS -f a -p 12345 -x 12345 -s {input.fasta} -m
 GTRGAMMA -# 100 -T 8 -n {output.fasta.name} -o
 {outgroup.sequence.names}
~~~

### Defining structurally variable haplotypes

To define the set of structurally distinct haplotypes for the evolutionary and biomedically important loci, we performed an all against all pairwise alignment for each of the haplotypes using the following minimap2 command (*99*):

~~~
 minimap2 -r 50000 -ax asm20 --eqx -Y
~~~

Sequences aligned to the same haplotype for at least 90% of their length at >99% identity without deletions or insertions of 50 kbp or more were grouped into a single structural haplotype and considered not structurally variable. Structurally variable haplotypes were then defined as the mutually exclusive groups where every haplotype in a given group did not align to the haplotype of any other group for >90% of its length at >99% identity. Similarly, haplotypes that were grouped together with GRCh38 or CHM13 were deemed to have “recapitulated the structural organization” of that particular reference.

### Variation graphs for SD loci

We applied minigraph v0.14 (*60*) to construct variation graphs using all structurally distinct haplotypes with the parameters:

~~~
 minigraph -xggs -L 5000 -r 100000 -t {threads} *.fasta
~~~

All haplotypes were aligned back to the graph to call variants:

~~~
 minigraph -x asm -t {threads} {input.gfa} {input.fasta}
~~~

### Methylation analysis

Methylation analysis was performed using the same data and methods described by Gershman et al., bioRxiv (*105*). In brief, CHM13 ultra-long ONT reads were aligned to the CHM13 reference with Winnowmap2 (*106*) with a k-mer size of 15 and filtered for primary alignments for read lengths greater than 50 kbp. To measure CpG methylation in nanopore data, we used Nanopolish (v0.13.2) (*64*) filtered methylation calls using the nanopore_methylation_utilities tool (https://github.com/timplab/nanopore-methylation-utilities), which uses a log-likelihood ratio of 1.5 as a threshold for calling methylation. Methylation data was then loaded into R for all downstream analysis with GenomicRanges (*107*) and dplyr (*108*).

### Custom ideogram and homology visualizations

Linear ideograms were constructed using the karyoploteR package (*109*) and circular ideograms were made using circlize (*110*). R code used to make these figures is shared for convenience at https://github.com/mrvollger/Vollger_2020_Figures; however, this is not a software package and is provided without extensive documentation or installation instructions.

Sequence homology plots were made with a modified version of Miropeats (*111*) that uses minimap2 to identify alignments. Code for the homology plots can be found in https://github.com/mrvollger/assembly_workflows under workflows/minimiro.smk (*112*–*114*). In brief, sequences are aligned using the following minimap2 parameters:

~~~
 minimap2 -x asm20 -r 200000 -s 100000 \
 -N 1000 --secondary=no\
 --cs {input.ref} {input.query} > {output.paf}
~~~

and then processed into a postscript file using scripts/minimiro.py and converted into a PDF.

## DATA AVAILABILITY

PacBio HiFi data has been deposited into NCBI Sequence Read Archive (SRA) under the following accessions: SRX7897688, SRX7897687, SRX7897686, and SRX7897685 for CHM13; SRR14407677 and SRR14407676 for CHM1; SRR10382244, SRR10382245, SRR10382248 and SRR10382249 for HG002; PRJNA540705 for NA12878; PRJEB36100 for HG00733 and HG00514; ERX4787609, ERX4787607, ERX4787606, ERX4782632, and ERX4781730 for NA19240; PRJNA701308 for HG01109, HG01243, HG02080, HG02723, HG03125, and HG03492; and PRJNA659034 and PRJNA691628 for all nonhuman primate samples. The complete T2T-CHM13 assembly and the CHM13 ONT data, including raw signal files (FAST5), base calls (FASTQ), and alignments (BAM/CRAM), are available at https://github.com/nanopore-wgs-consortium/chm13. The assembly can also be found on NCBI (GCA_009914755.2) and all the read data has been uploaded on SRA under the BioProject identifier PRJNA686988. Two human PacBio Iso-Seq datasets from fetal brain and testis are accessioned under NCBI BioProject PRJNA659539. The canonical rDNA unit used to estimate copy number can be found on the NCBI nucleotide repository (KY962518.1). Human and nonhuman primate genome assemblies, SD annotations, methylation data, and Liftoff gene models can be found on Zenodo (DOI: 10.5281/zenodo.4721956).

## ACKNOWLEDGEMENTS

The authors thank T. Brown for help in editing this manuscript. This work was supported, in part, by the Intramural Research Program of the National Human Genome Research Institute, National Institutes of Health (S.N., S.K., and A.M.P.), grants from the U.S. National Institutes of Health (NIH grants 5R01HG002385 to E.E.E.; 5U01HG010971 to E.E.E.; 1U01HG010973 to E.E.E.; 1R01HG011274 to K.H.M.; 5R01HG009190 to W.T.; and U41HG007234 to M.D.), and a grant from Futuro in Ricerca (2010-RBFR103CE3 to M.V.). E.E.E. is an investigator of the Howard Hughes Medical Institute.

## AUTHOR CONTRIBUTIONS

Identification of SDs in T2T-CHM13 and analysis: M.R.V.; PacBio genome sequence generation: K.M.M, A.M.L., K.H.; FISH experiments and analysis; L.M., M.V., M.R.V., E.E.E.; Iso-Seq analysis: P.C.D., M.R.V., R.L.; *TBC1D3* analysis: X.G., M.R.V.; copy number analysis: M.R.V., W.T.H.; inversion analysis: D.P., M.R.V.; T2T-CHM13 assembly generation: S.N., S.K., A.M.P.; refinement of SD annotations near centromeres K.H.M., M.R.V.; UCSC browser: M.D., W.T.H., M.R.V.; methylation analysis: M.R.V., A.G., W.T., E.E.E.; analysis of regions with genomic instability: M.R.V., A.S.; organization of tables: M.R.V., P.C.D., X.G.; organization of supplementary material: M.R.V.; display items: M.R.V., X.G., P.C.D.; manuscript writing: M.R.V., E.E.E. X.G. with input from all authors.

## Notes

https://doi.org/10.5281/zenodo.4721956

### Competing Interest Statement

The authors have declared no competing interest.

