## Supplemental material for "Segmental duplications and their variation in a complete human genome"

### Supplemental Notes

**Supplemental note 1:**
We report 81.3 Mbp of SD sequence that overlaps with the new or structurally variable sequence in T2T-CHM13 assembly. This differs from the 68.3 Mbp reported by Nurk et al. (*1*) because we considered all SDs that overlapped with new sequence whereas Nurk et al. uses a strict intersection. For example, if a 75 kbp SD overlapped 50 kbp of new sequence we would report 75 kbp of new SD sequence whereas Nurk et al. would report 50 kbp.

**Supplemental note 2:**
Nurk et al. reports 140 new protein-coding genes (*1*), and we report 182 new genes with multiple exons and an open reading frame (ORF). This difference comes from different gene annotation sets and different filtering steps. Nurk et al. uses CAT (*2*) to annotate genes in the assembly which they then supplement with Liftoff (*3*) while considering only new gene copies that are 100% identical to previously annotated paralogs. We used Liftoff to generate our gene annotations and allowed new paralogs to diverge by up to 15% from previously annotated paralogs if they still had an ORF— and this difference accounts for the additional genes we identify. Of the 140 genes, 58 overlap with the 182 we report. This difference comes from an additional set of filters we used to offset the false positives that were identified by allowing 15% divergence. Our additional filters were that the gene must have an ORF of at least 200 bp and have multiple exons.

### Supplemental Figures

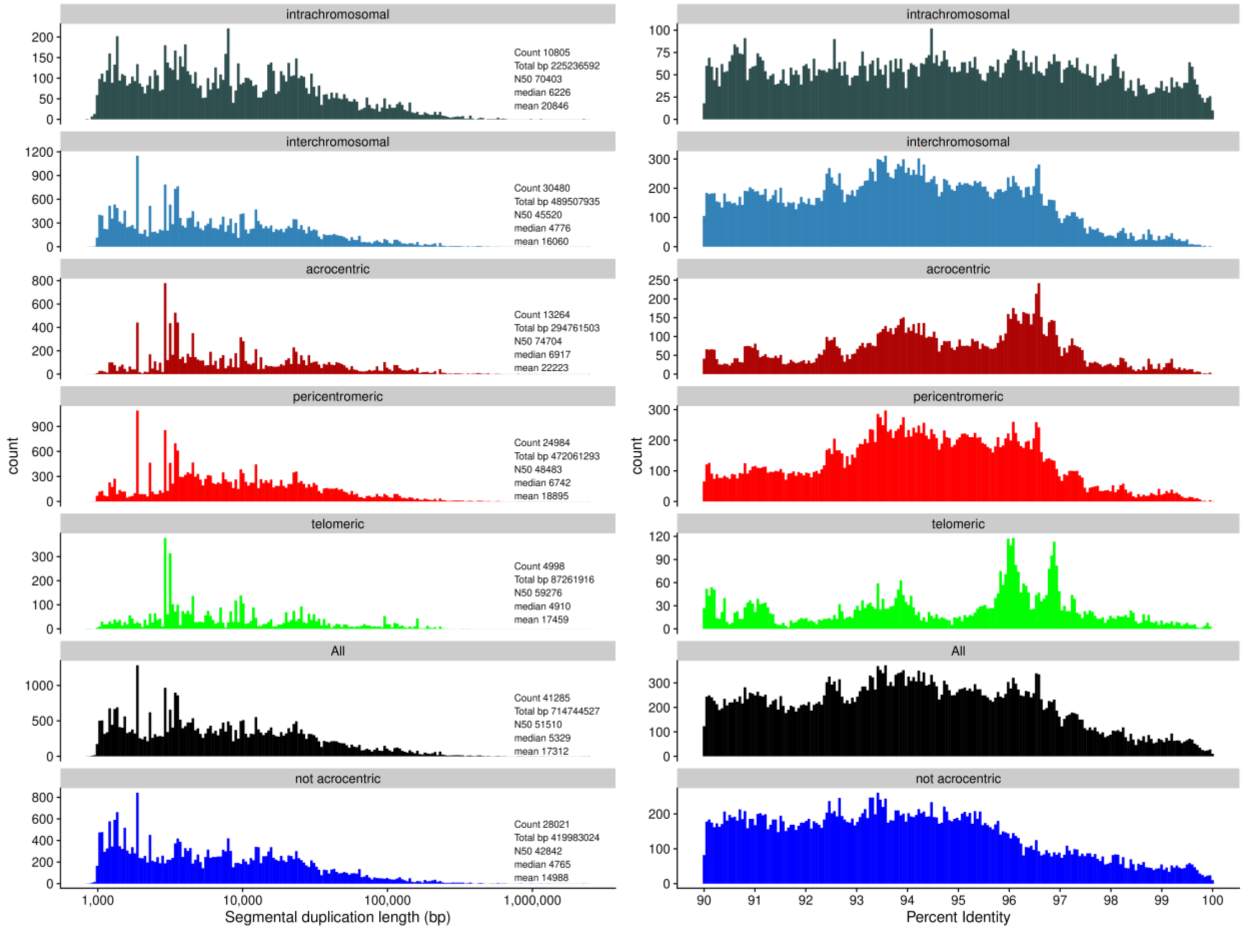


**Figure S1. Comparison of SD length and identity in different regions of the genome.** The length (left) and identity (right) of SDs across commonly delineated regions of the genome (colors).

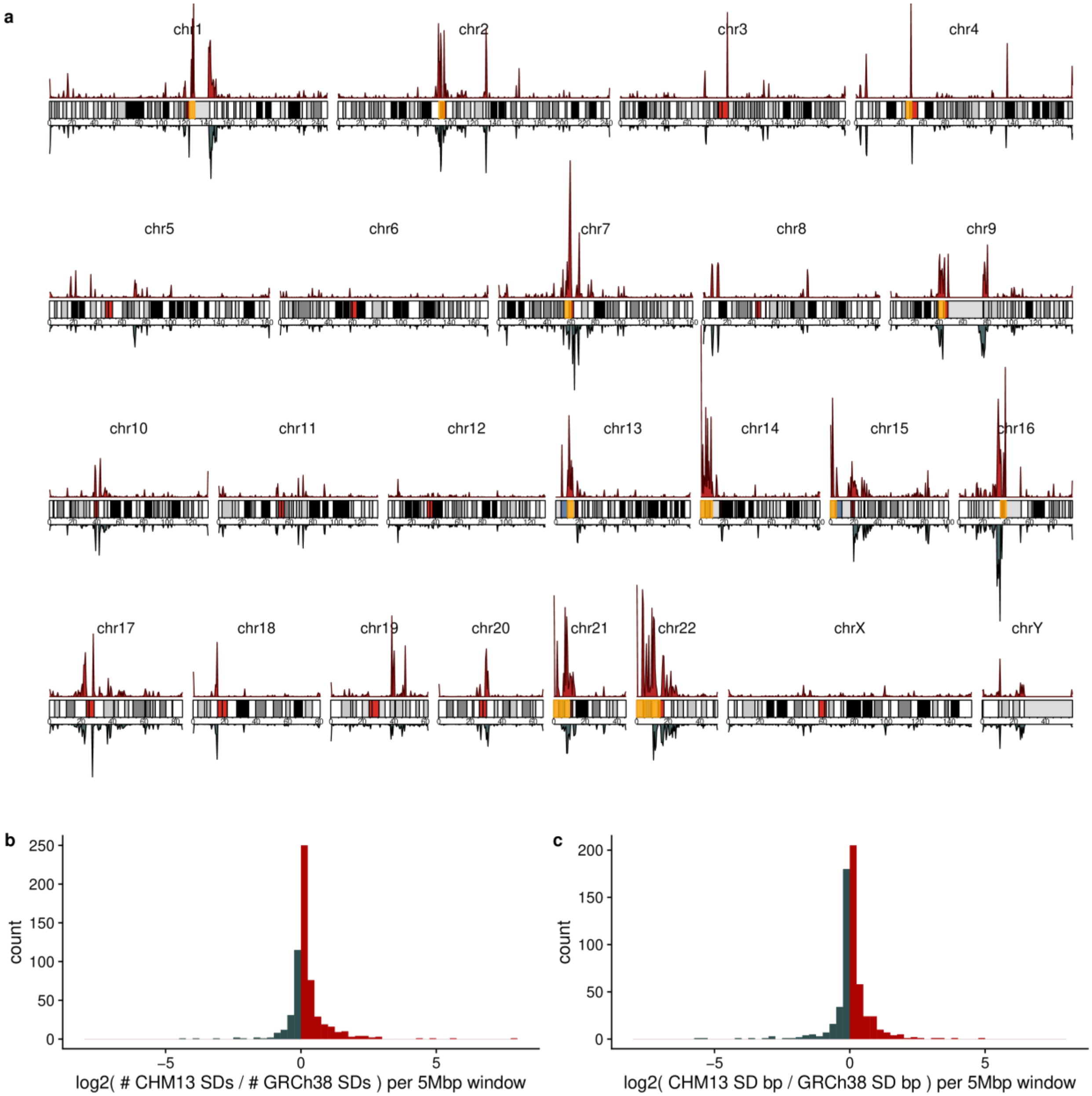


**Figure S2. SD density comparison between T2T-CHM13 and GRCh38.**
**a)** Density of SDs in T2T-CHM13 (red) and GRCh38 (blue). In the ideogram highlighted in orange are the 15 regions with the largest increase in the number of SDs. **b)** Histogram showing the $\log_2$ fold change between the number of SDs in T2T-CHM13 and GRCh38 per non-overlapping 5 Mbp window. **c)** Histogram showing the $\log_2$ fold change between the number of bp in SDs for T2T-CHM13 and GRCh38 per non-overlapping 5 Mbp window.

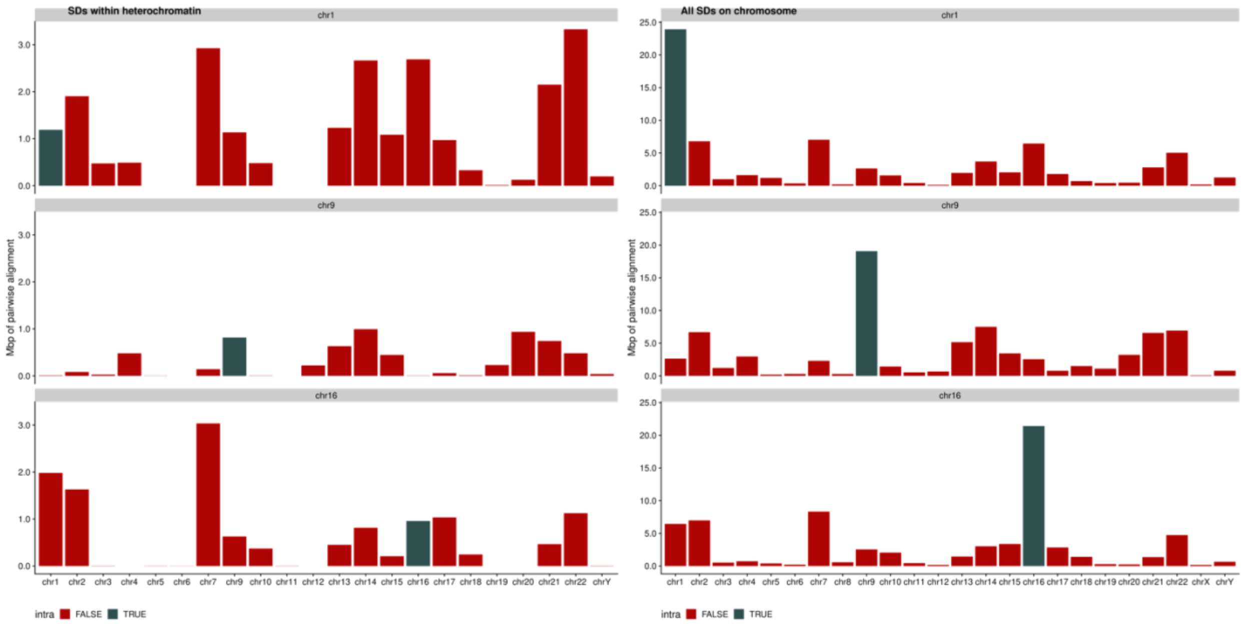


**Figure S3. SDs within heterochromatin on chromosomes 1, 9, and 16.**
This figure shows where the SDs that separate the HSAT and centromere arrays on chromosomes 1, 9, and 16 align to (left) compared to the overall distribution of that chromosome (right). Blue are intrachromosomal SDs and red are interchromosomal.

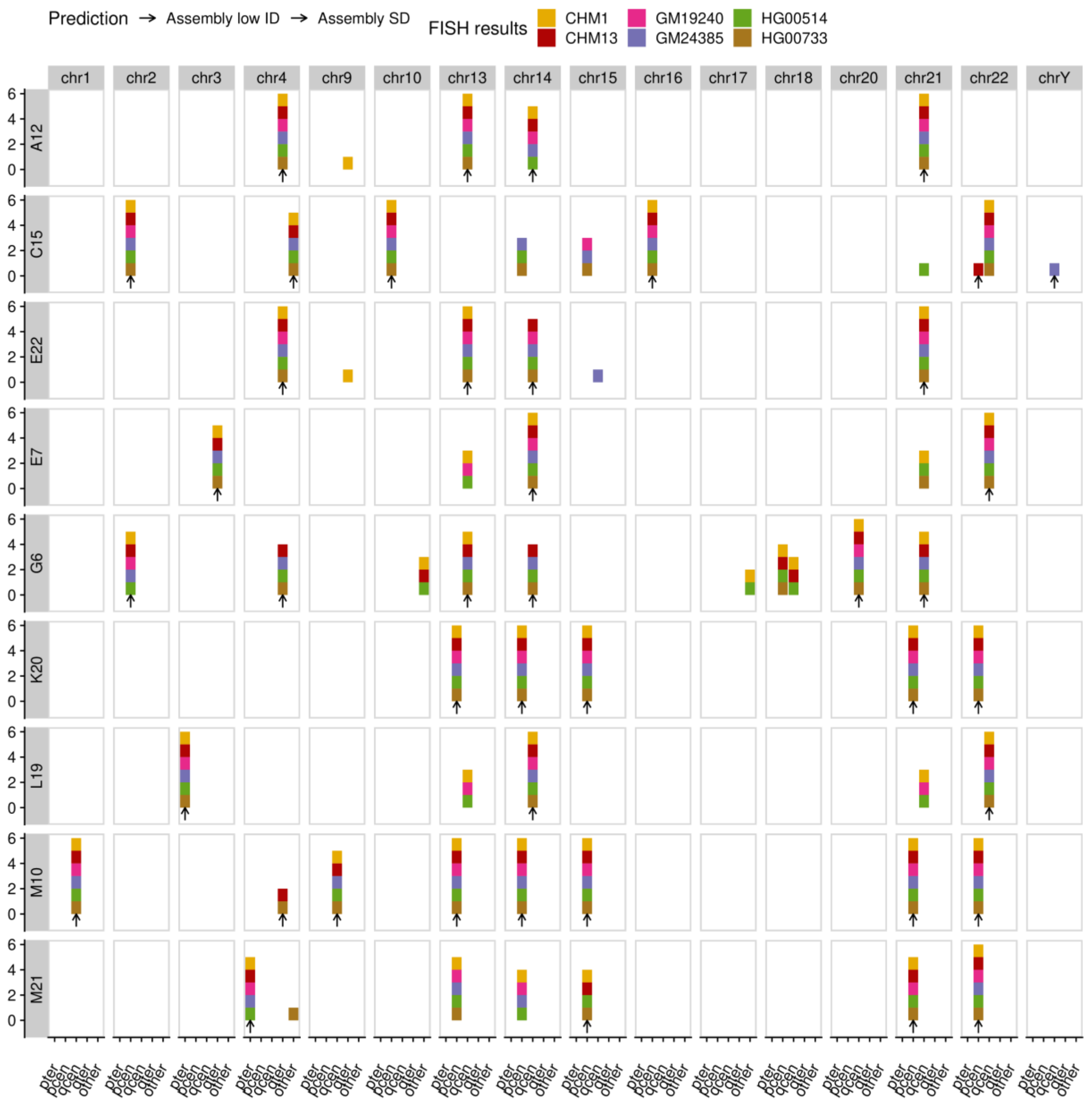


**Figure S4. FISH support for new CHM13 duplications.**
This table shows the location and chromosomes of FISH signals (x) for each probe (y) across the different cell lines (color). Black shows the predicted locations of FISH signals from the T2T-CHM13 assembly.

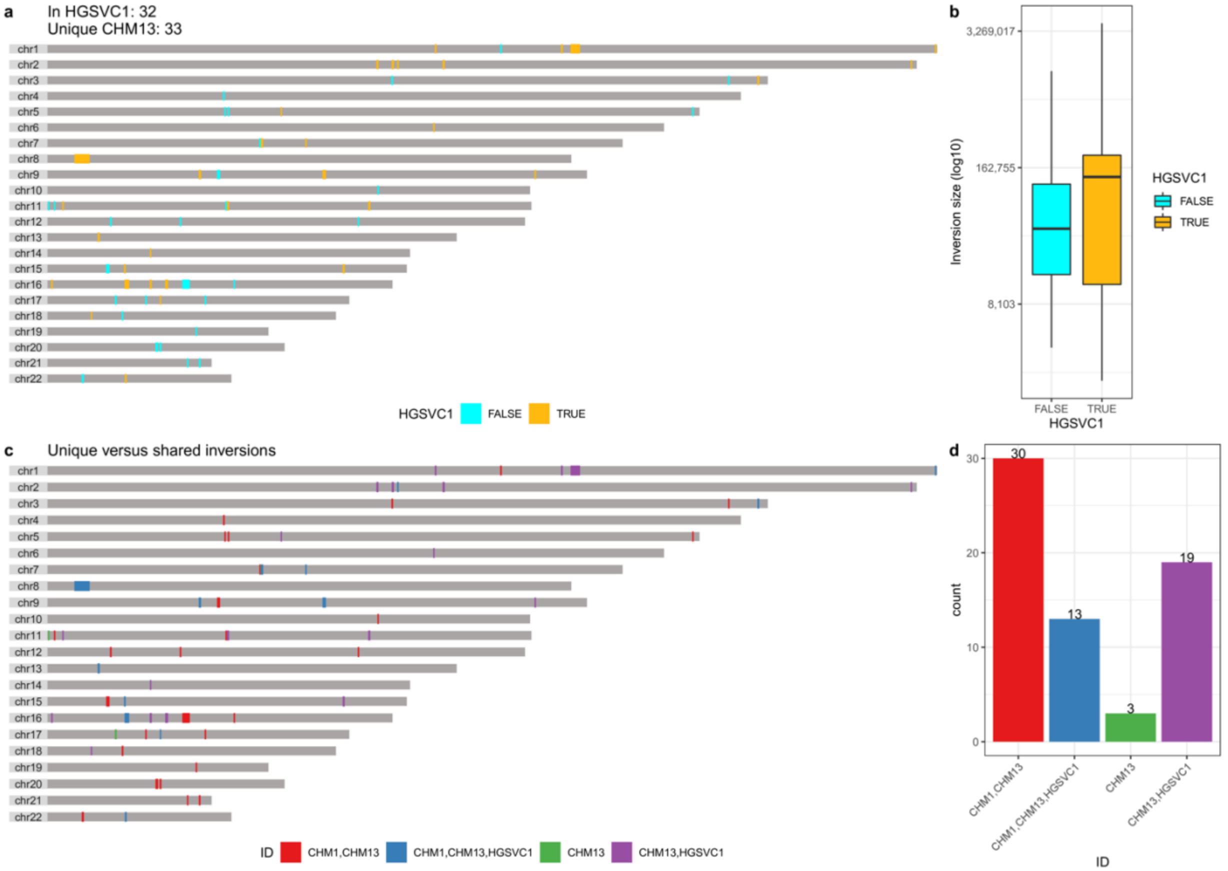


**Figure S5. CHM13 inversions supported by Strand-seq.**

**a)** Locations of inversions in CHM13 relative to GRCh38 as identified with Strand-seq. The color indicates whether the inversion is shared (orange) with at least one sample from HGSVC1 (*4*) or unique to CHM13 (cyan). **b)** Size distribution of inversions in CHM13. **c)** Comparison of inversions shared between CHM1 and CHM13. d) Bar chart showing the counts of shared and unique inversions in CHM13.

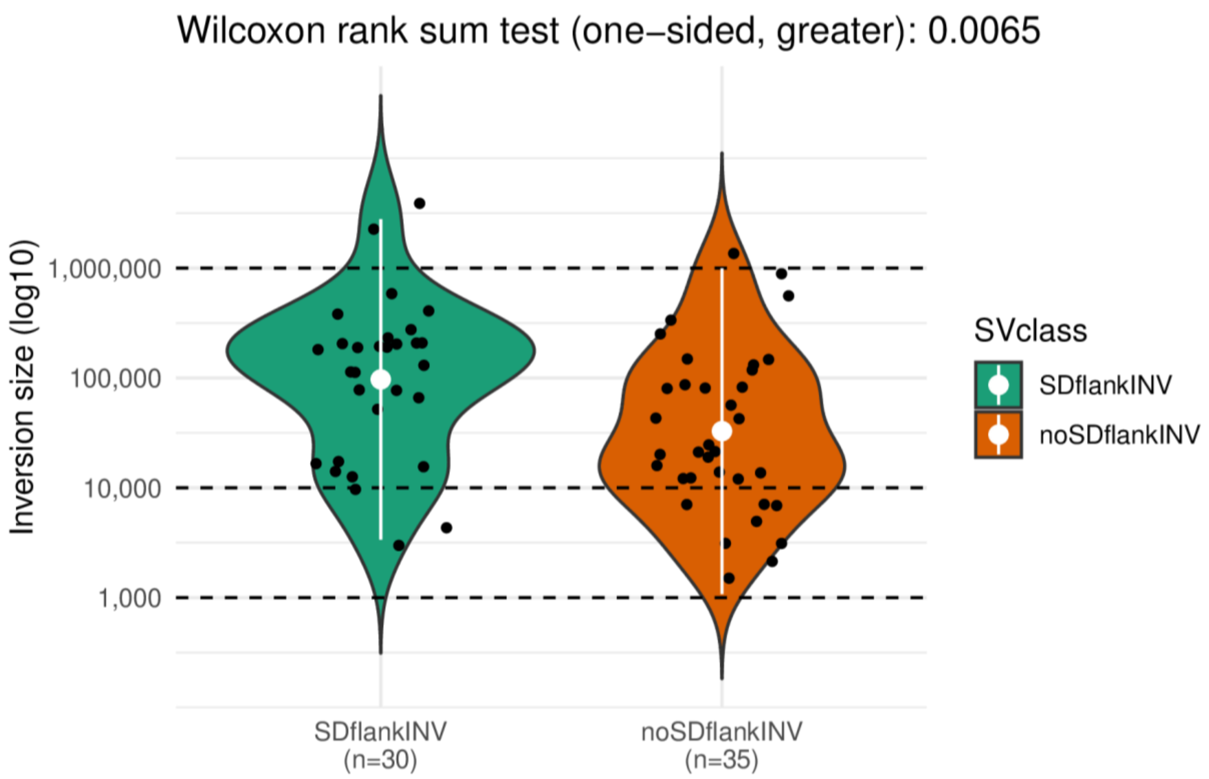


**Figure S6. Length of CHM13 inversions.**
The length of inversions in CHM13 as predicted by Strand-seq stratified by the presence of flanking SDs (green) or lack thereof (orange). Inversions flanked by SDs are significantly longer than other inversions ($p = 0.0065$, one-sided Wilcoxon rank-sum test).

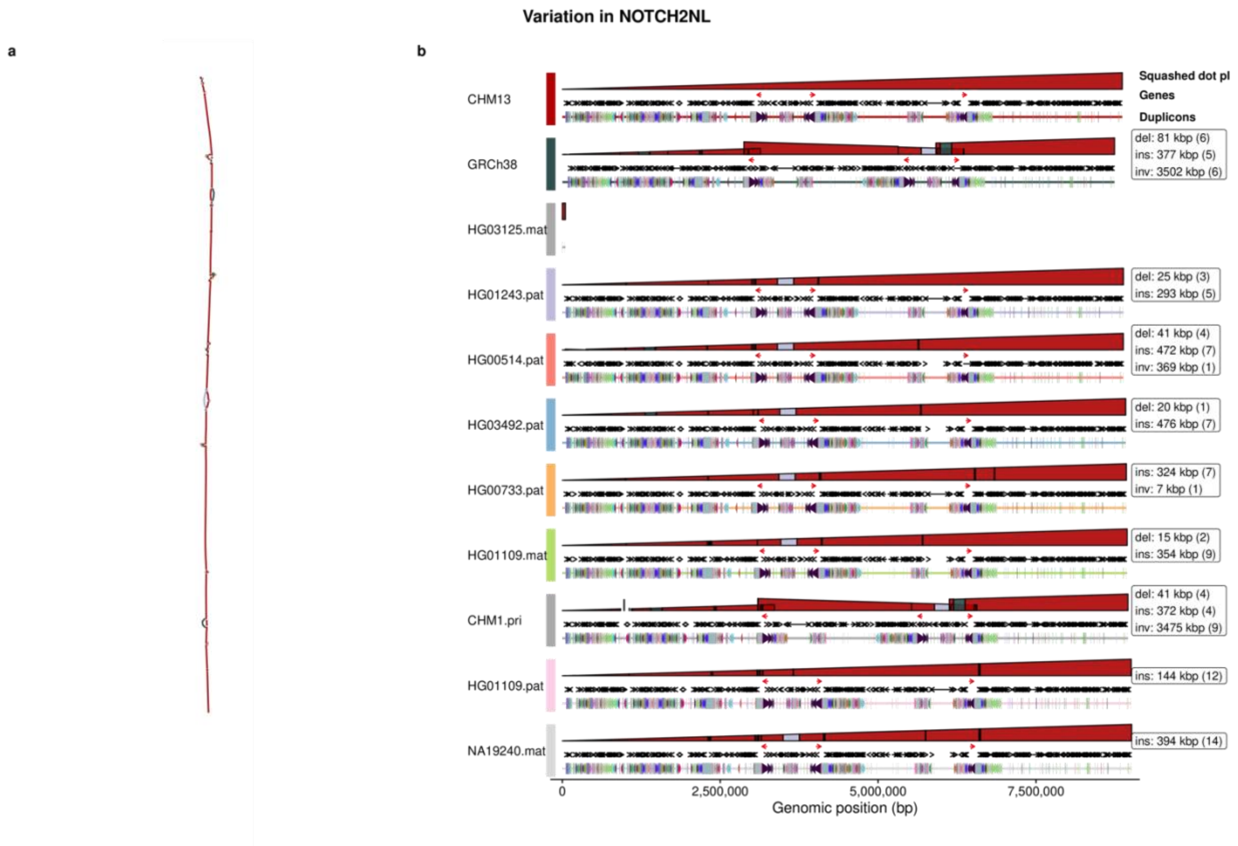


**Figure S7. Pangenome graph of *NOTCH2NL* and *SRGAP2*.**

**a)** Variation in human haplotypes across the *NOTCH2NL* expansion site: a graph representation [rGFA generated using minigraph (*5*)] of the locus where colors indicate the source genome for the sequence. The graph visualization was created using the software tool Bandage (*6*). **b)** The path for each haplotype-resolved assembly through the graph. The “squashed dot plot” represents a vertically compressed dot plot comparing the haplotype-resolved sequence (horizontal) against the graph (vertical). Color represents the source haplotype for the vertical sequence. Structural variants can be identified from discontinuities in height (deletion), changes between colors (insertion), or changes in the direction of the polygon (inversion *NOTCH2NL*, the gene of interest, is shown with red arrows and other genic content in the region are shown with black arrows. The final line is a duplicon track showing the ancestral duplications (color) that make up the larger duplication block.

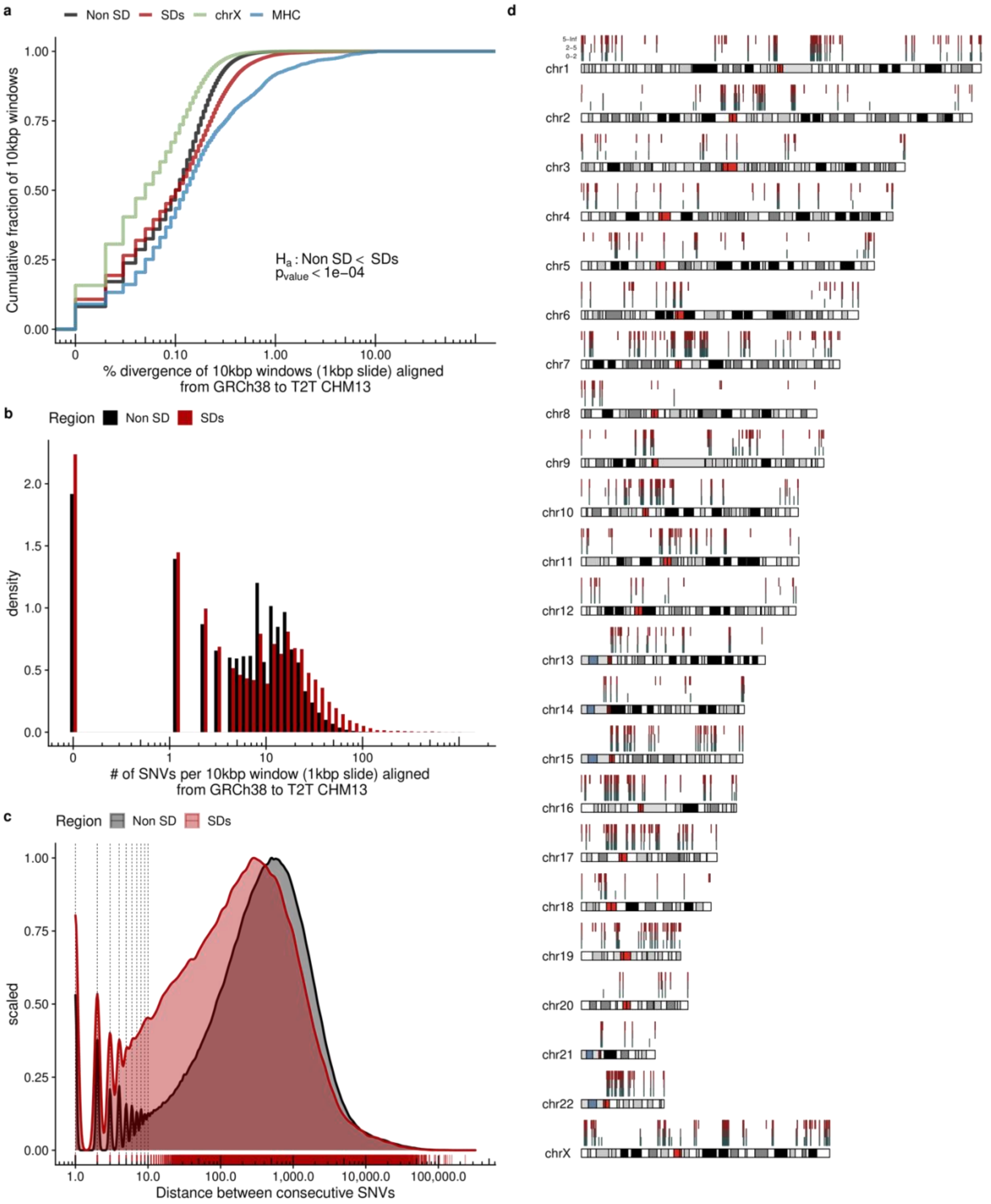


**Figure S8. Single-nucleotide variants (SNVs) in SDs between T2T-CHM13 and GRCh38.**
a) Divergence of 10 kbp windows with synteny between GRCh38 and T2T-CHM13.
b) Distribution of the number of SNVs per 10 kbp windows aligned from GRCh38 to T2T-CHM13 in unique and SD regions. c) Distribution of the distance between SNVs in the syntenic regions of GRCh38 and T2T-CHM13. d) SD regions with synteny between T2T-CHM13 and GRCh38

and their average levels of single-nucleotide variation in 1 kbp windows. The bottom row has windows of SD with 0-2 SNVs per kbp, middle row 2-5 SNVs per kbp, and top row is greater than 5 SNVs per kbp.

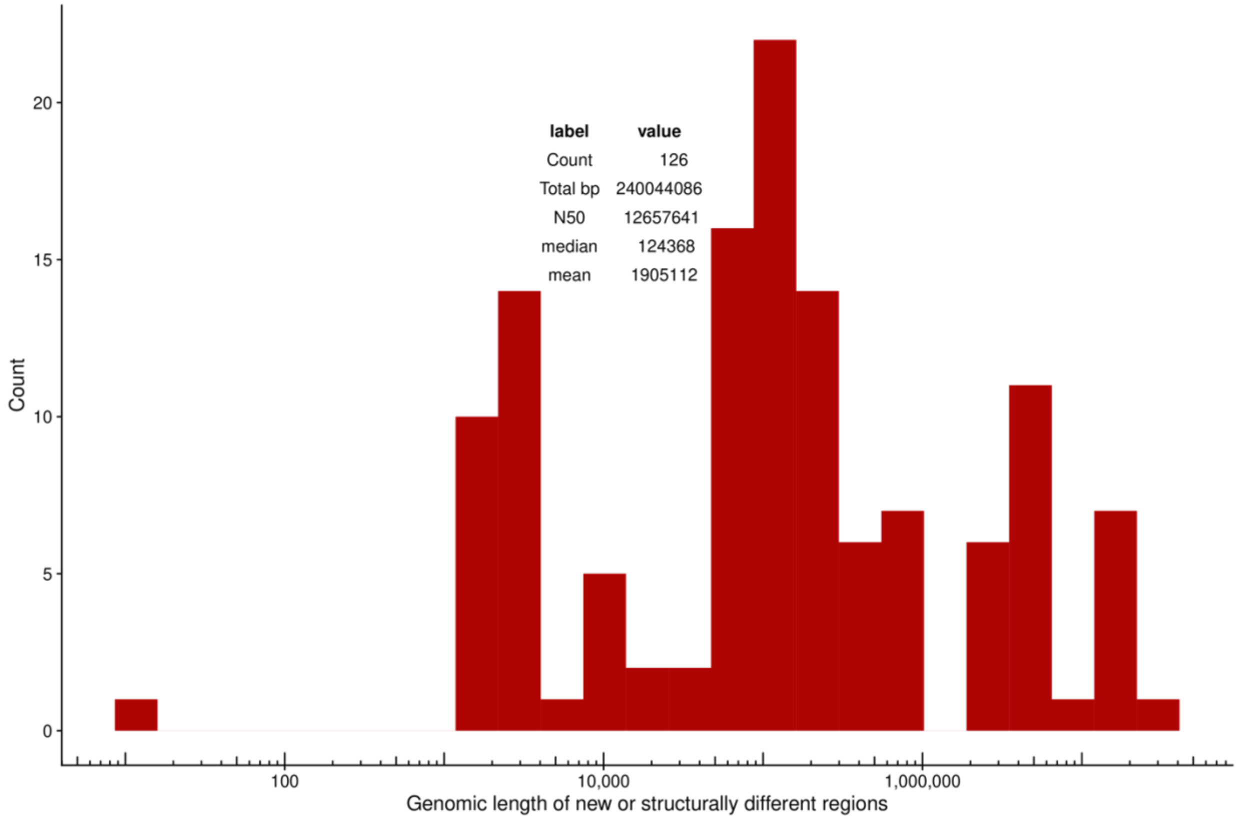


**Figure S9. Size distribution of non-syntenic regions between GRCh38 and T2T-CHM13.** Histogram showing the size of non-syntenic regions (Methods) between GRCh38 and T2T-CHM13, and a table of statistics on the lengths of the region.

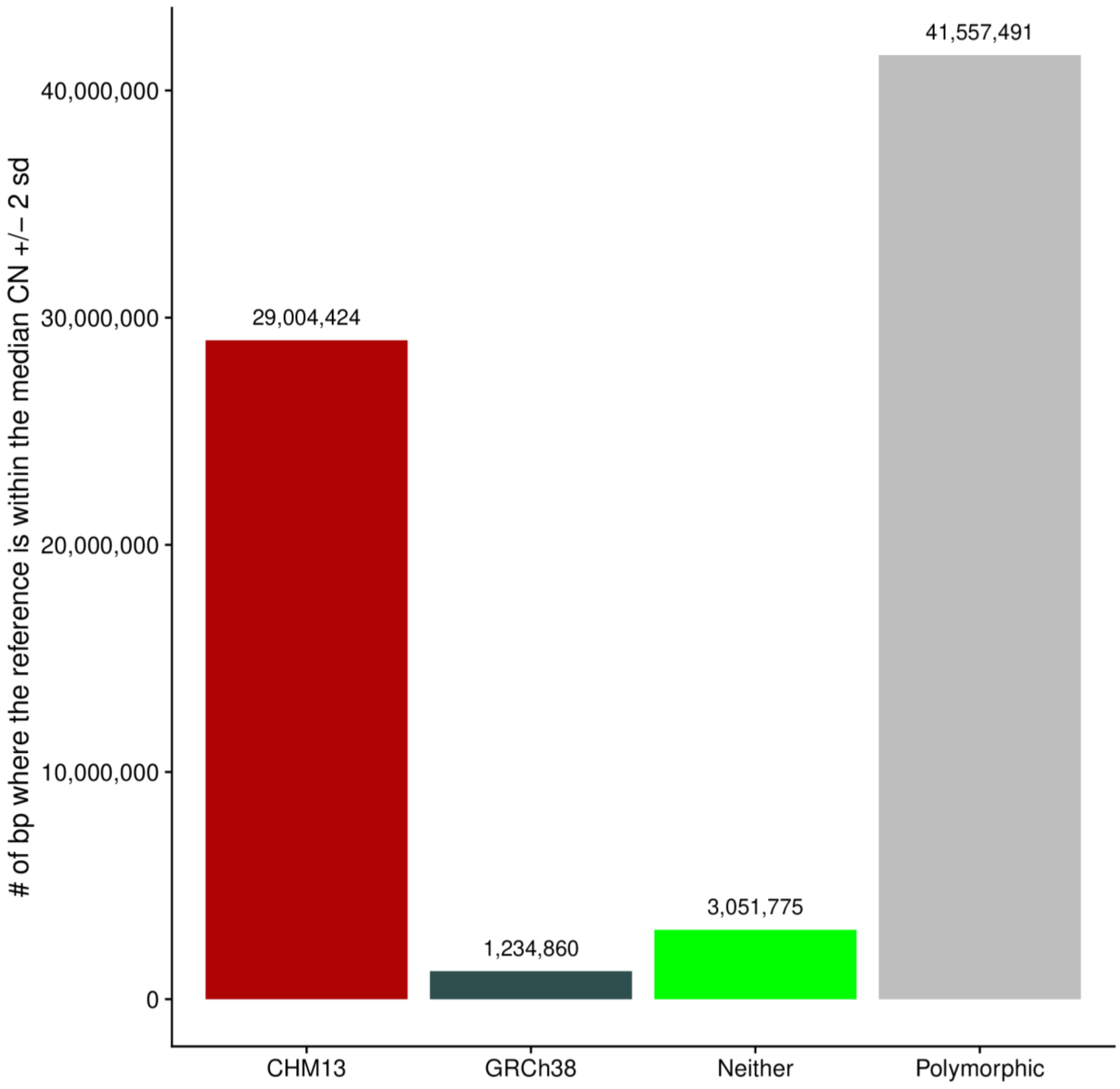


**Figure S10. Non-syntenic regions where the reference copy number reflects SGDP.** Copy number (CN) of SD regions that are new or structurally different in T2T-CHM13 compared to GRCh38 and 268 individuals from the SGDP (*7*). The histogram shows the number in Mbp where the median sample CN from SGDP was within two standard deviations (s.d.) of the given assembly [T2T-CHM13 (red), GRCh38 (blue), neither (green), or both (equal CN)].

a

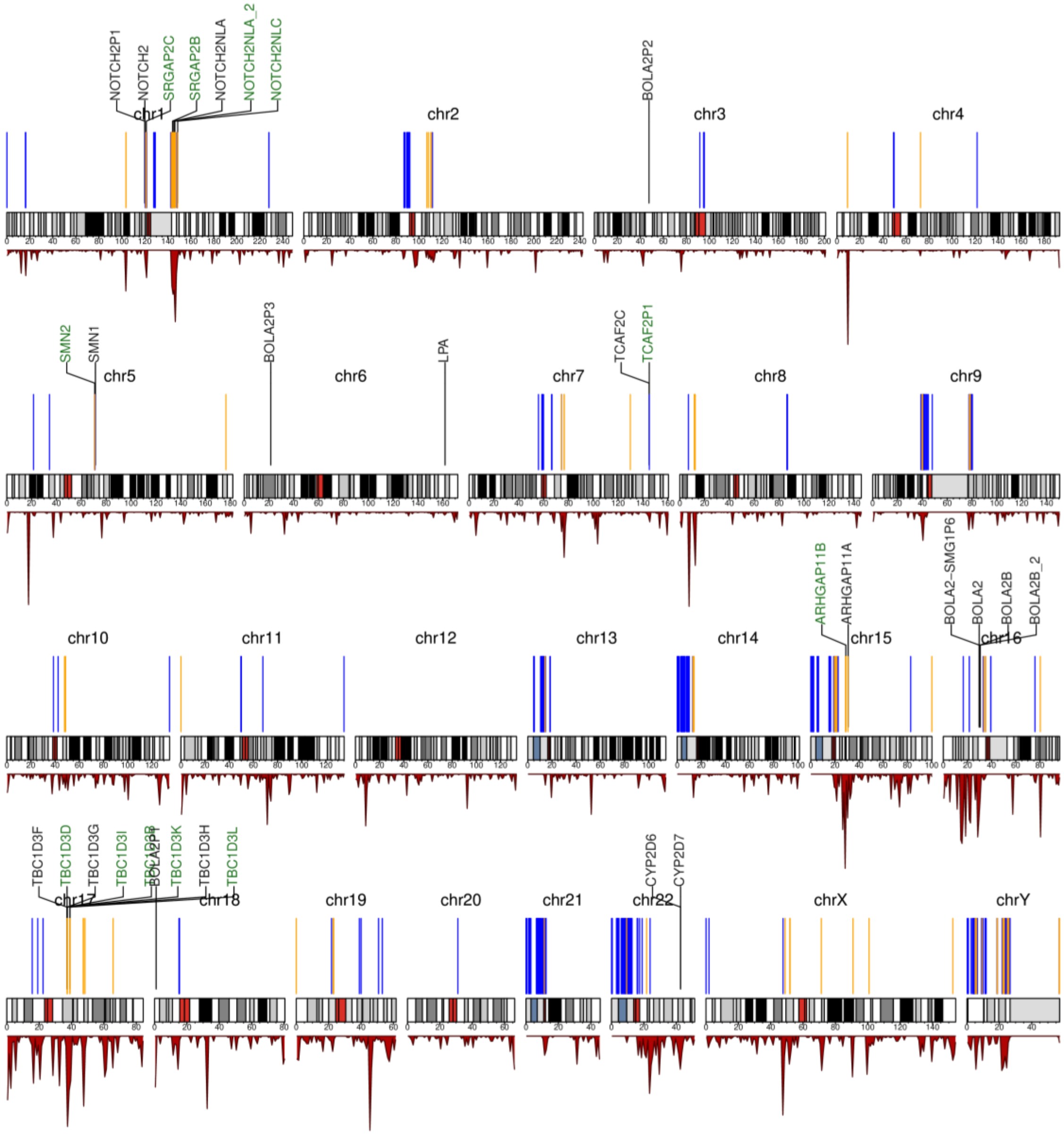


**Figure S11. Genic SD expansions in T2T-CHM13 relative to chimpanzee.**
The blue (no genes) and orange (containing genes) peaks in the ideogram show regions of expansion in CHM13 relative to the Clint_PTR assembly within SD space. The bottom panel shows the density of genic SDs in T2T-CHM13. The genes highlighted as biomedically or evolutionarily important loci are labeled and colored green if they are part of a human expansion.

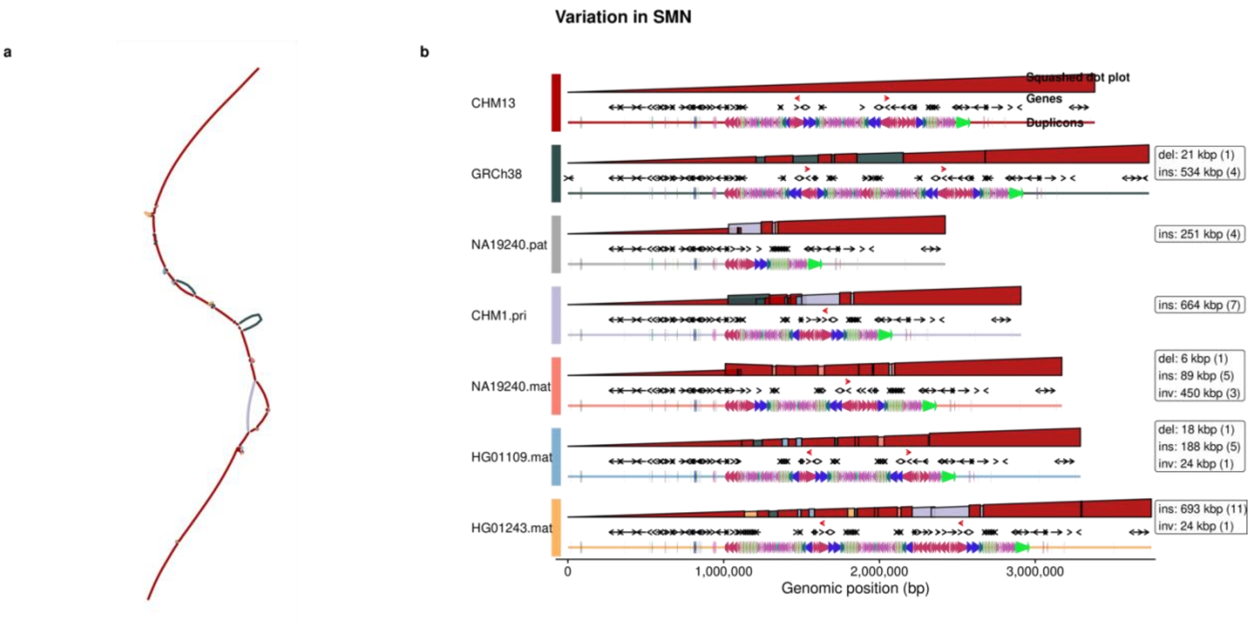


**Figure S12. Pangenome graph of *SMN*.**
For a description of the elements within this figure, see Figure S7.

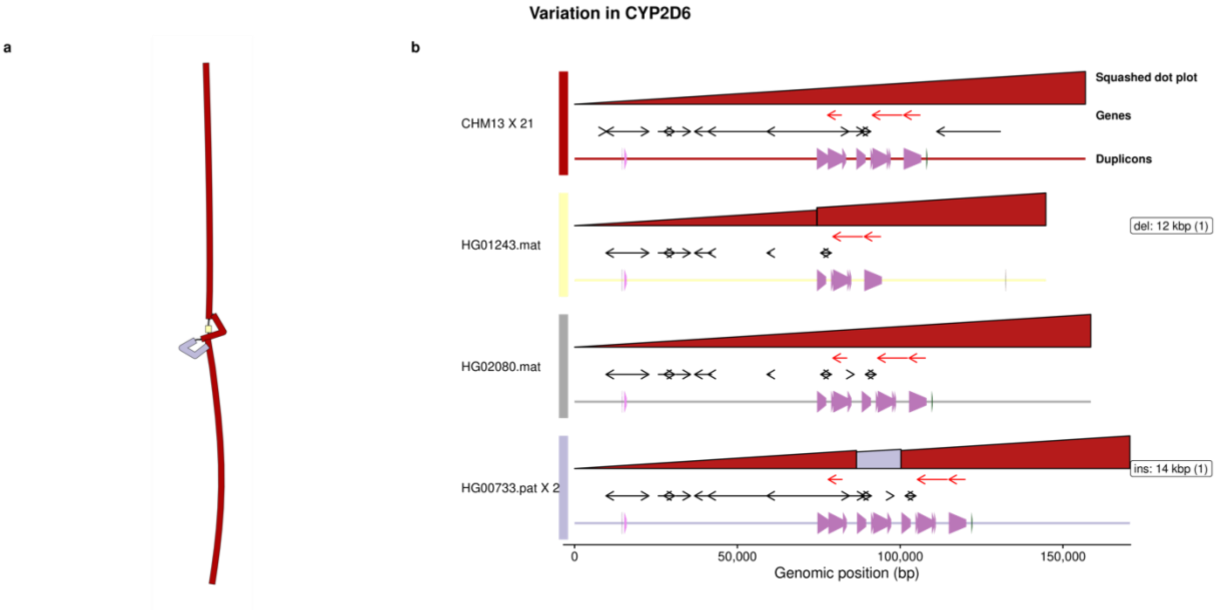


**Figure S13. Pangenome graph of *CYP2D6*.**
For a description of the elements within this figure, see Figure S7.

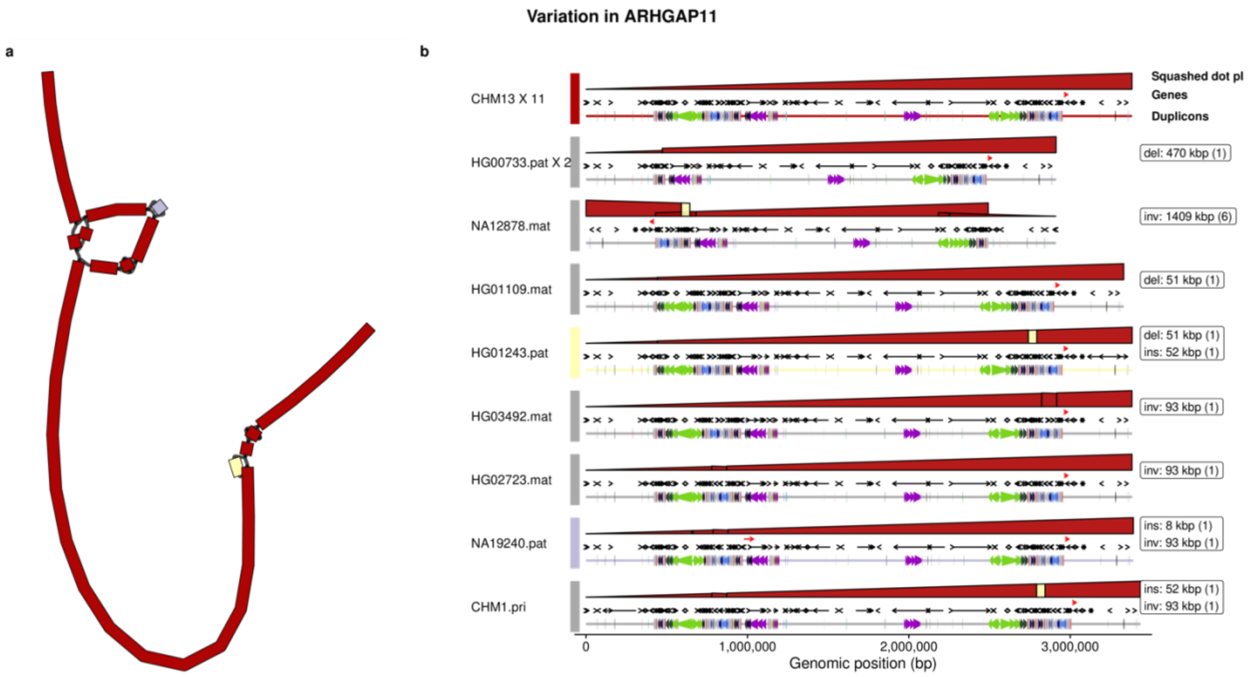


**Figure S14. Pangenome graph of *ARHGAP11*.**
For a description of the elements within this figure, see Figure S7.

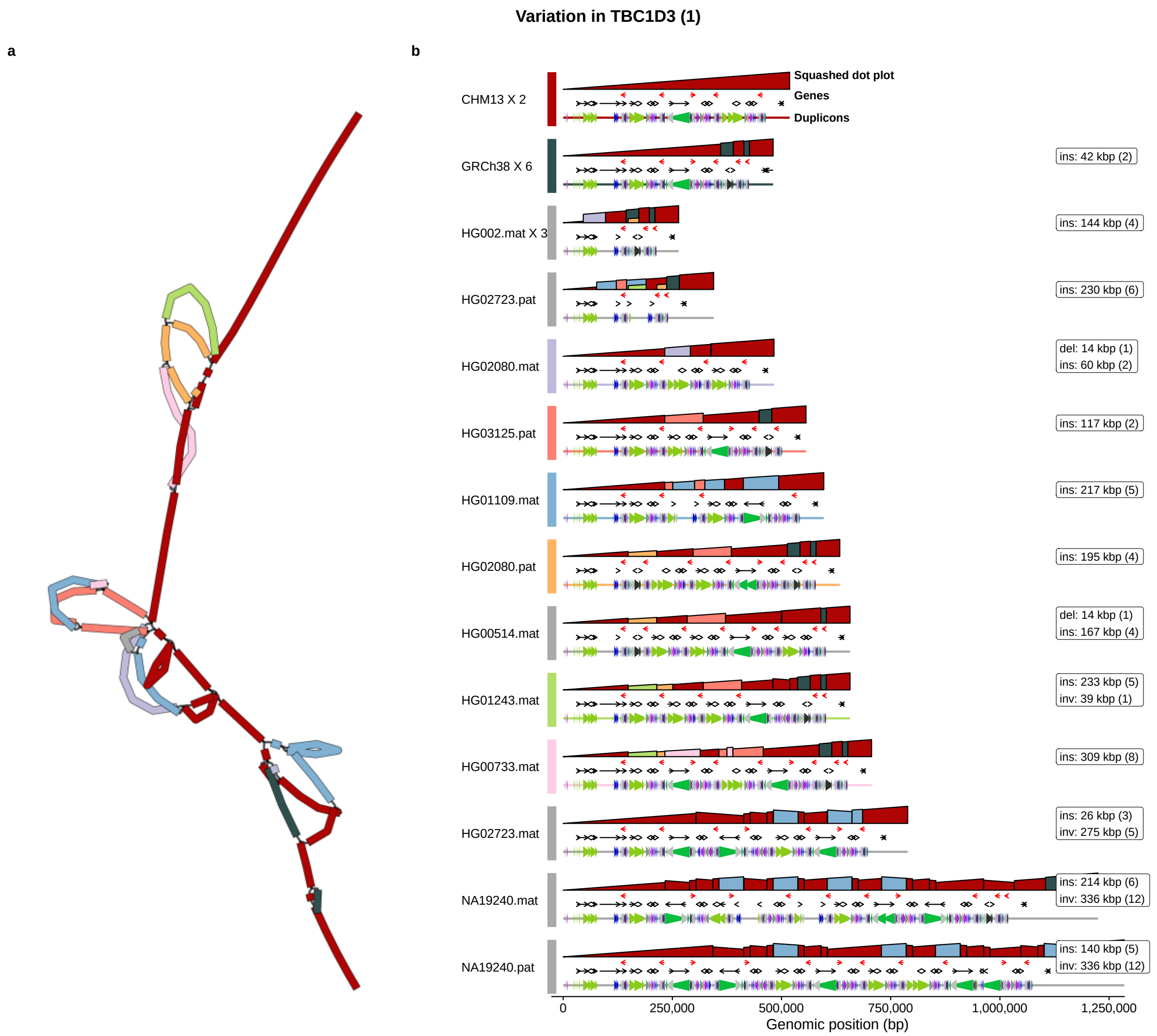


**Figure S15. Pangenome graph of *TBC1D3* expansion site one.**
For a description of the elements within this figure, see Figure S7.

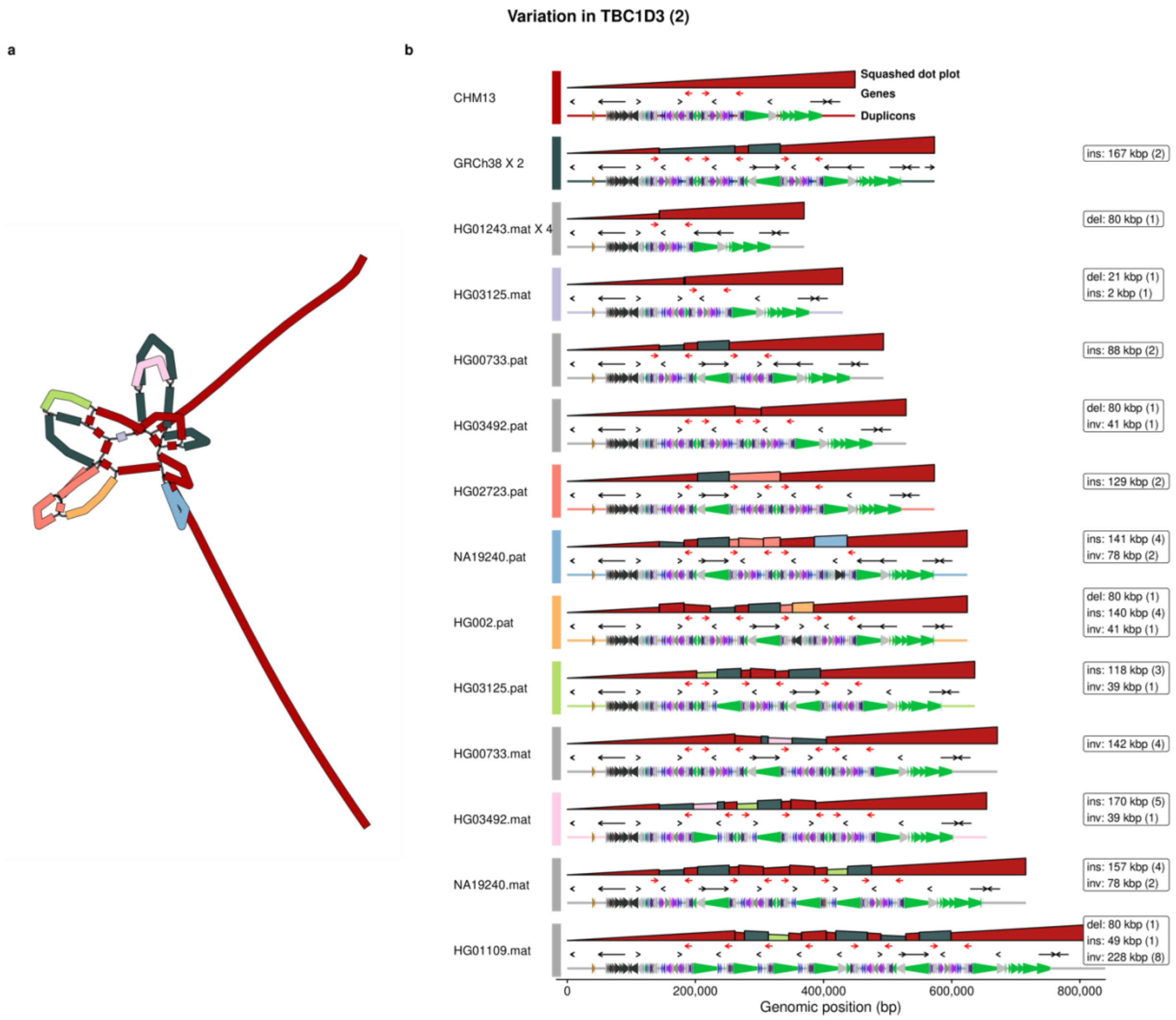


**Figure S16. Pangenome graph of *TBC1D3* expansion site two.**
For a description of the elements within this figure, see Figure S7.

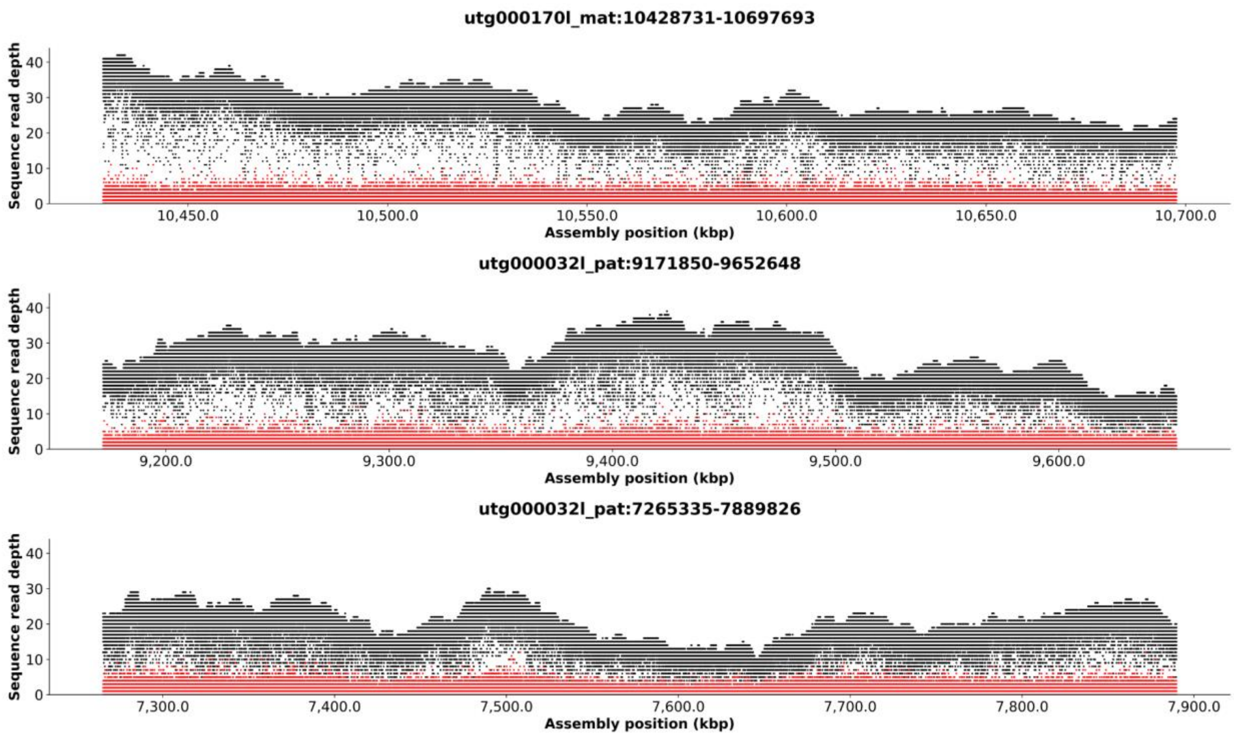


**Figure S17. Validation of assembly using ONT coverage over *TBC1D3* for HG002.**
Ultra-long ONT coverage of HG002 across the maternal haplotype of *TBC1D3* expansion site one, and the coverage across the maternal and paternal haplotypes of *TBC1D3* expansion site two. Black dots show the coverage of the most frequent base at each genomic position and red dots show the coverage of the second most frequent base.

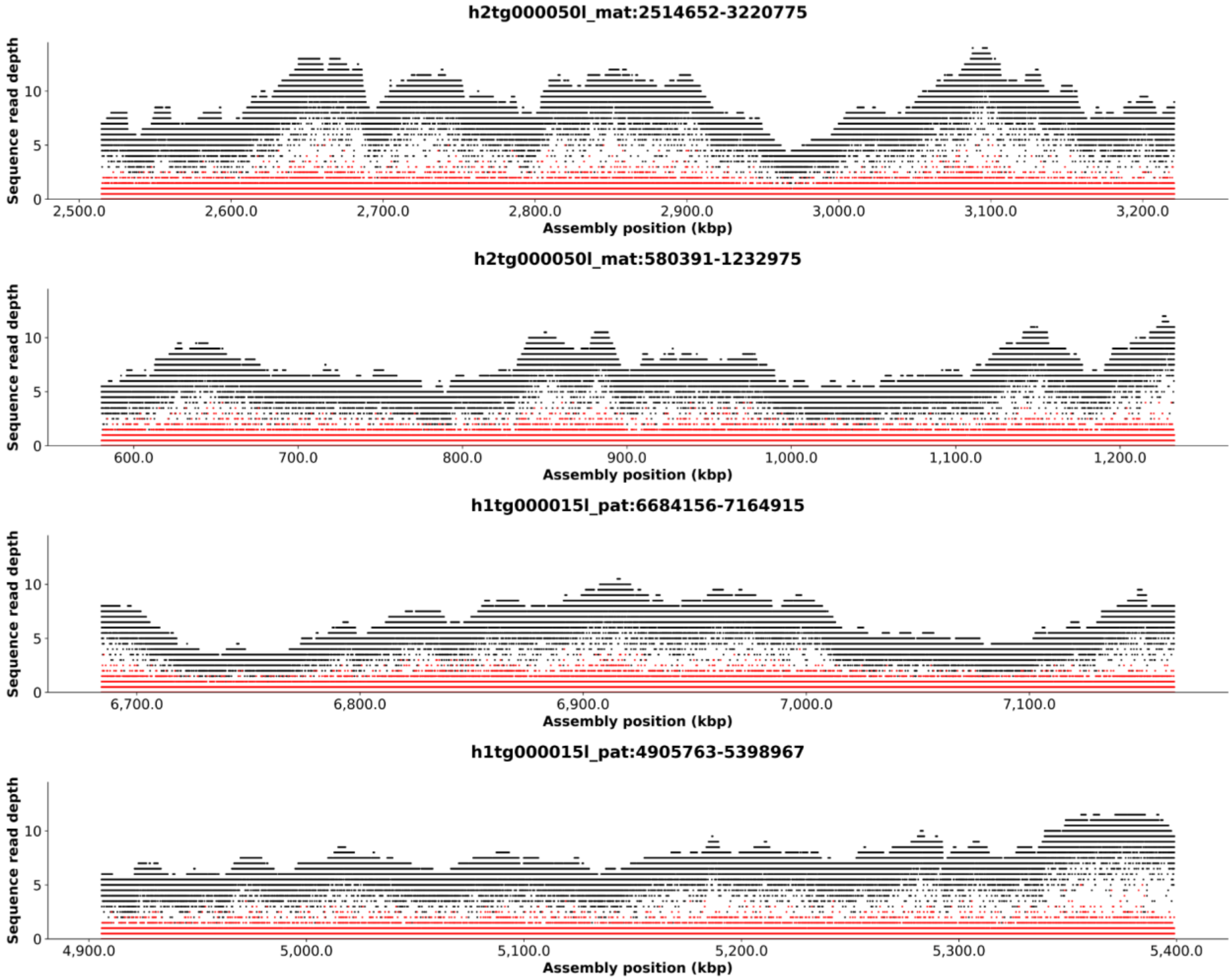


**Figure S18. Validation of assembly using ONT coverage over *TBC1D3* for HG00733.** Ultra-long ONT coverage of HG00733 across the maternal and paternal haplotypes of *TBC1D3* expansion site one, and the coverage across the maternal and paternal haplotypes of *TBC1D3* expansion site two. Black dots show the coverage of the most frequent base at each genomic position and red dots show the coverage of the second most frequent base.

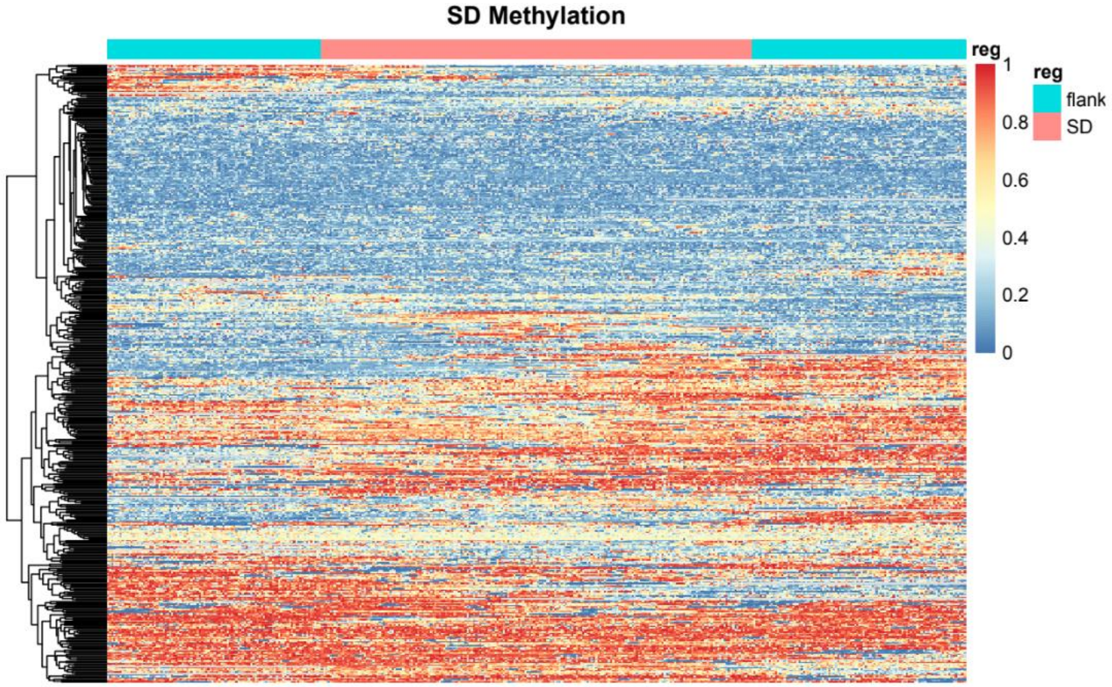


**Figure S19. Clustering of methylation status in SD blocks.**
Heatmap of CpG methylation of all SD blocks with at least 50 kbp of flanking sequence clustered using the “pheatmap” package in R. The horizontal annotation shows in cyan the 50 kbp of unique flanking sequence and red the SD block.

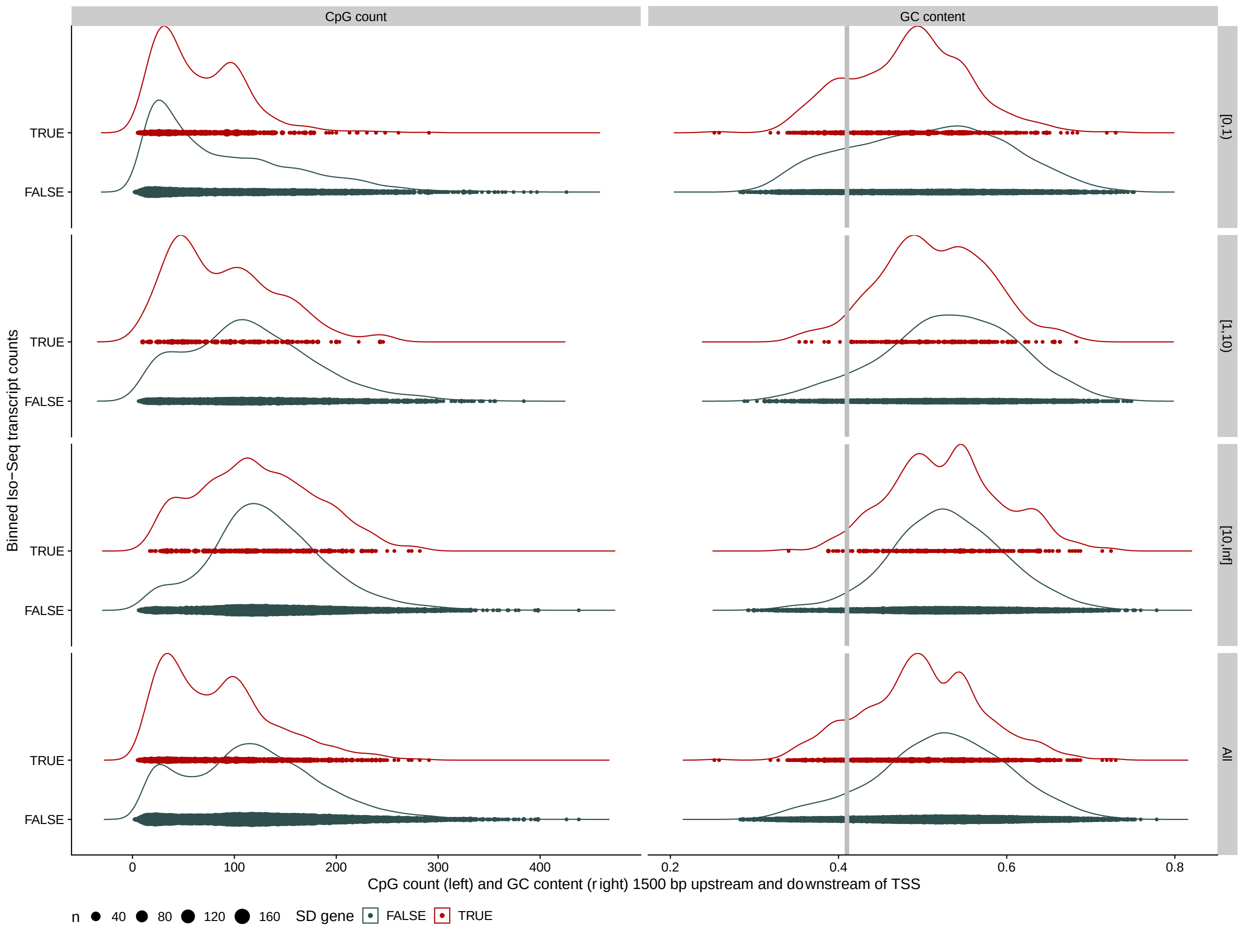


**Figure S20. CpG and GC content within 1,500 bp of the transcription start site (TSS).** Shown are the density of the number of CpGs within +/-1,500 bp of the TSS (left), and the density of bases that are G or C within +/-1,500 bp of the TSS (right), both stratified by the level of Iso-Seq transcription (vertical positioning) and SD content (color).

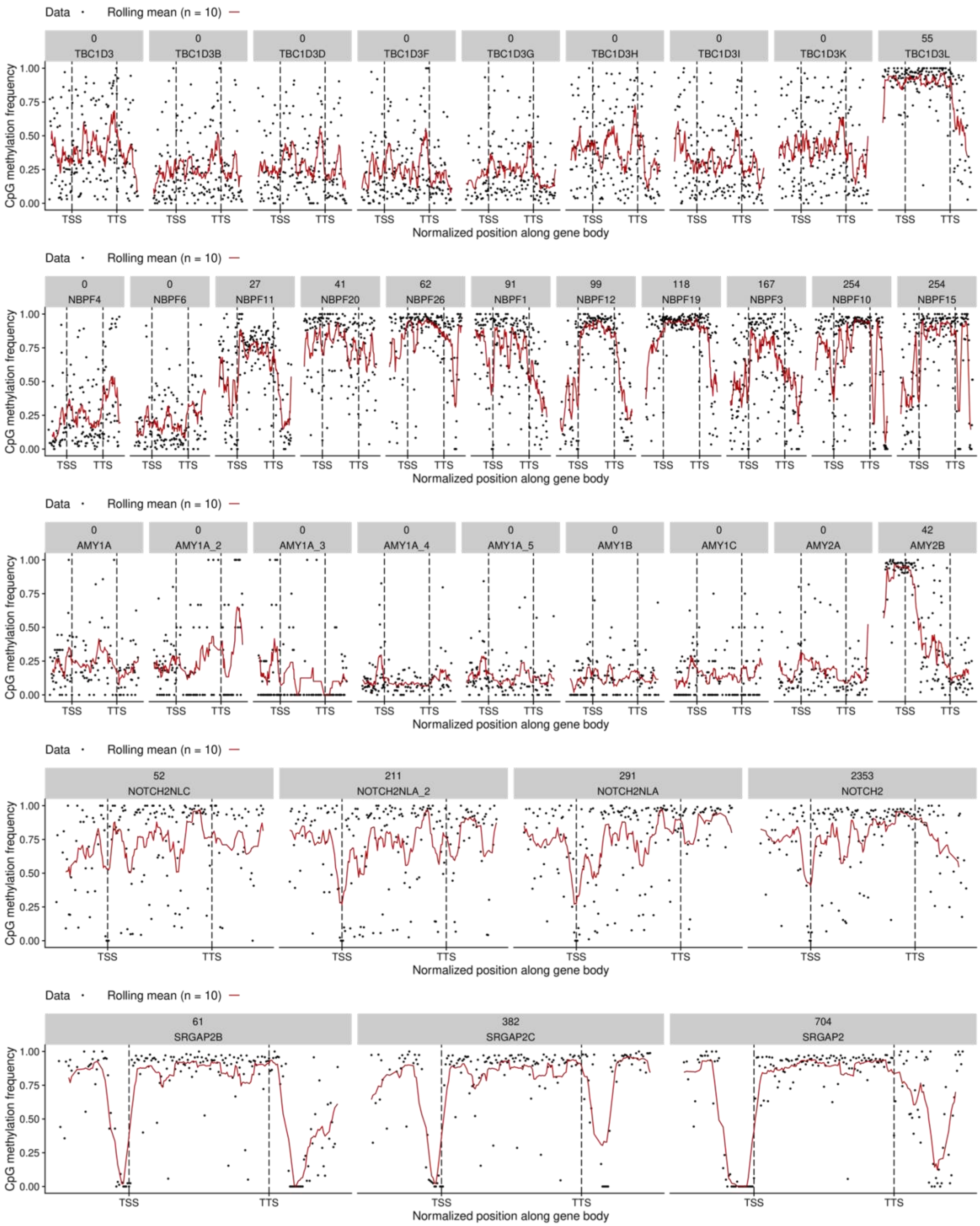


**Figure S21. Methylation and transcription levels across multi-copy gene families.**

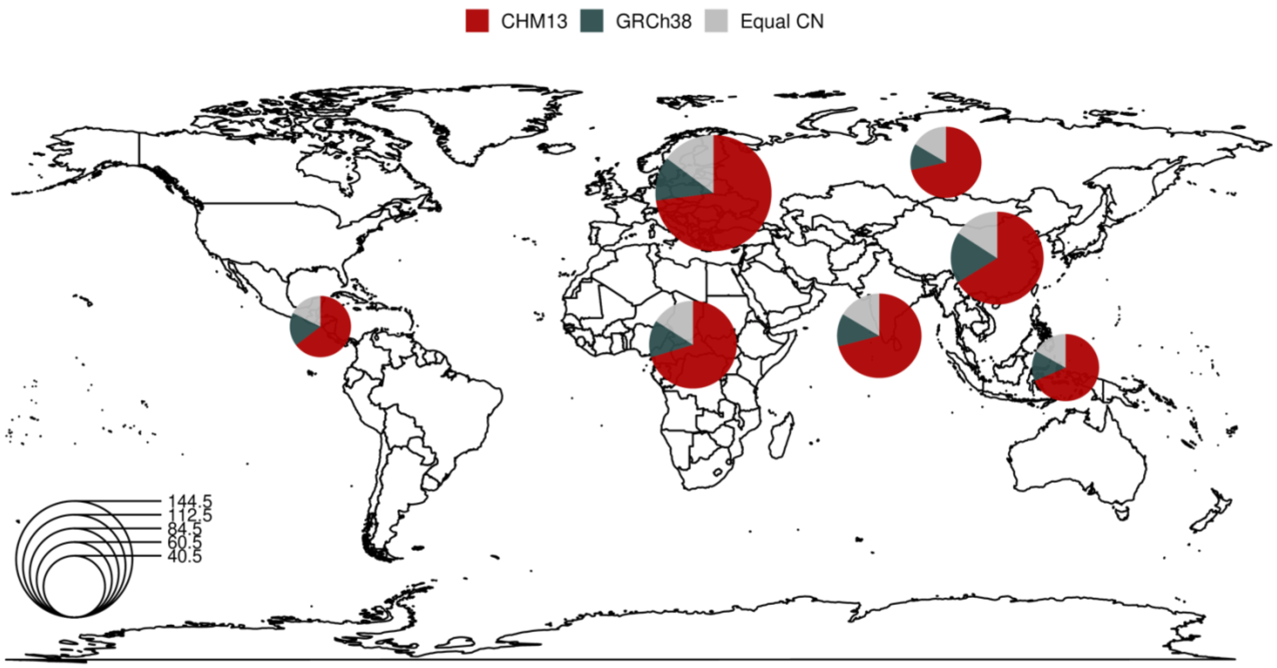


**Figure S22. Better CN representation of the SGDP across super populations.**
This figure shows pie charts for each human super population from the Simon Genome Diversity Panel (SGDP, n = 268). Individual pie charts show the relative fraction of SGDP samples for every non-syntenic SD region where the individual's CN more closely matches the T2T-CHM13 CN (red), GRCh38 CN (blue), or is represented equally well by both (gray).
